# Fine-mapping of nuclear compartments using ultra-deep Hi-C shows that active promoter and enhancer elements localize in the active A compartment even when adjacent sequences do not

**DOI:** 10.1101/2021.10.03.462599

**Authors:** Huiya Gu, Hannah Harris, Moshe Olshansky, Yossi Eliaz, Akshay Krishna, Achyuth Kalluchi, Mozes Jacobs, Gesine Cauer, Melanie Pham, Suhas S.P. Rao, Olga Dudchenko, Arina Omer, Kiana Mohajeri, Sungjae Kim, Michael H Nichols, Eric S. Davis, Devika Udupa, Aviva Presser Aiden, Victor G. Corces, Douglas H. Phanstiel, William Stafford Noble, Jeong-Sun Seo, Michael E. Talkowski, Erez Lieberman Aiden, M. Jordan Rowley

## Abstract

Megabase-scale intervals of active, gene-rich and inactive, gene-poor chromatin are known to segregate, forming the A and B compartments. Fine mapping of the contents of these A and B compartments has been hitherto impossible, owing to the extraordinary sequencing depths required to distinguish between the long-range contact patterns of individual loci, and to the computational complexity of the associated calculations. Here, we generate the largest published *in situ* Hi-C map to date, spanning 33 billion contacts. We also develop a computational method, dubbed PCA of Sparse, SUper Massive Matrices (POSSUMM), that is capable of efficiently calculating eigenvectors for sparse matrices with millions of rows and columns. Applying POSSUMM to our Hi-C dataset makes it possible to assign loci to the A and B compartment at 500 bp resolution. We find that loci frequently alternate between compartments as one moves along the contour of the genome, such that the median compartment interval is only 12.5 kb long. Contrary to the findings in coarse-resolution compartment profiles, we find that individual genes are not uniformly positioned in either the A compartment or the B compartment. Instead, essentially all (95%) active gene promoters localize in the A compartment, but the likelihood of localizing in the A compartment declines along the body of active genes, such that the transcriptional termini of long genes (>60 kb) tend to localize in the B compartment. Similarly, nearly all active enhancers elements (95%) localize in the A compartment, even when the flanking sequences are comprised entirely of inactive chromatin and localize in the B compartment. These results are consistent with a model in which DNA-bound regulatory complexes give rise to phase separation at the scale of individual DNA elements.

## Main

The nucleus of the human genome is partitioned into distinct spatial compartments, such that stretches of active chromatin tend to lie in one compartment, called the A compartment, and stretches of inactive chromatin tends to lie in the other, called the B compartment^1^. Compartmentalization was first identified using Hi-C, a method that relies on DNA-DNA proximity ligation to create maps reflecting the spatial arrangement of the genome^1^. Loci in the same spatial compartment exhibit relatively frequent contacts in a Hi-C map, even when they lie far apart along a chromosome, or on entirely different chromosomes^1,2^. Accurate classification of the resulting genome-wide contact patterns requires a large number of contacts to be characterized at each locus. As such, genome-wide compartment profiles have only been generated, in the past, at resolutions ranging from 40 kb – 1 Mb^1–3^. Moreover, extant compartment detection algorithms require operations, such as calculation of principal eigenvectors^1^, which are computationally intractable when the underlying matrices have millions of rows and columns – such as high-resolution Hi-C matrices.

Although the compartments as a whole are often thought to form as a consequence of phase separation^3–6^, the low resolution of compartment profiles has made it difficult to determine the protein mechanisms that underlie this process.

Here, we construct an *in situ* Hi-C map in lymphoblastoid cells spanning 42 billion read-pairs and 33 billion contacts. This map contains an average of 22,000 contacts for every kilobase of genome sequence. We combine this map with a novel algorithm, dubbed POSSUMM, which greatly accelerates the calculation of the principal eigenvector and the largest eigenvalues of a massive, sparse matrix. This makes it possible to, e.g., calculate the principal eigenvector for correlation matrices containing millions of rows, and billions of nonzero entries. Combining our ultra-deep map with POSSUMM, we find that it is possible to map the contents of the A and B compartments with 500 bp resolution, a 100-fold improvement in resolution. We also show that when we classify loops based on their appearance, at fine resolution, in our ultra-deep map, it becomes possible to distinguish between loops that form by extrusion and those that form via non-extrusion mechanisms.

### Generation of an ultra-deep in situ Hi-C map in lymphoblastoid cells spanning 33 billion contacts

We produced an ultra-deep Hi-C map using lymphoblastoid cells from a panel of 17 individuals, obtaining over 42 billion PE150 read-pairs. This map was generated by aggregating the results of over 150 individual Hi-C experiments. In order to enhance the resolution of the maps, we used a variety of 4-cutter restriction enzymes in the different experiments, thus enhancing the density of cut sites across the genome. Together, these experiments yielded 33 billion contacts after alignment, deduplication, and quality filtering (Table S1). The resulting dataset is far deeper than any prior published Hi-C map. By comparison, the average published Hi-C map contains roughly 300 million contacts; 93% of Hi-C maps in the 4DNucleome database^7^ have less than 1 billion contacts (Fig. S1A, Table S2); and the widely used lymphoblastoid Hi-C map generated in Rao et al. contains 4.9 billion contacts (Fig. 1A).

**Figure 1.**
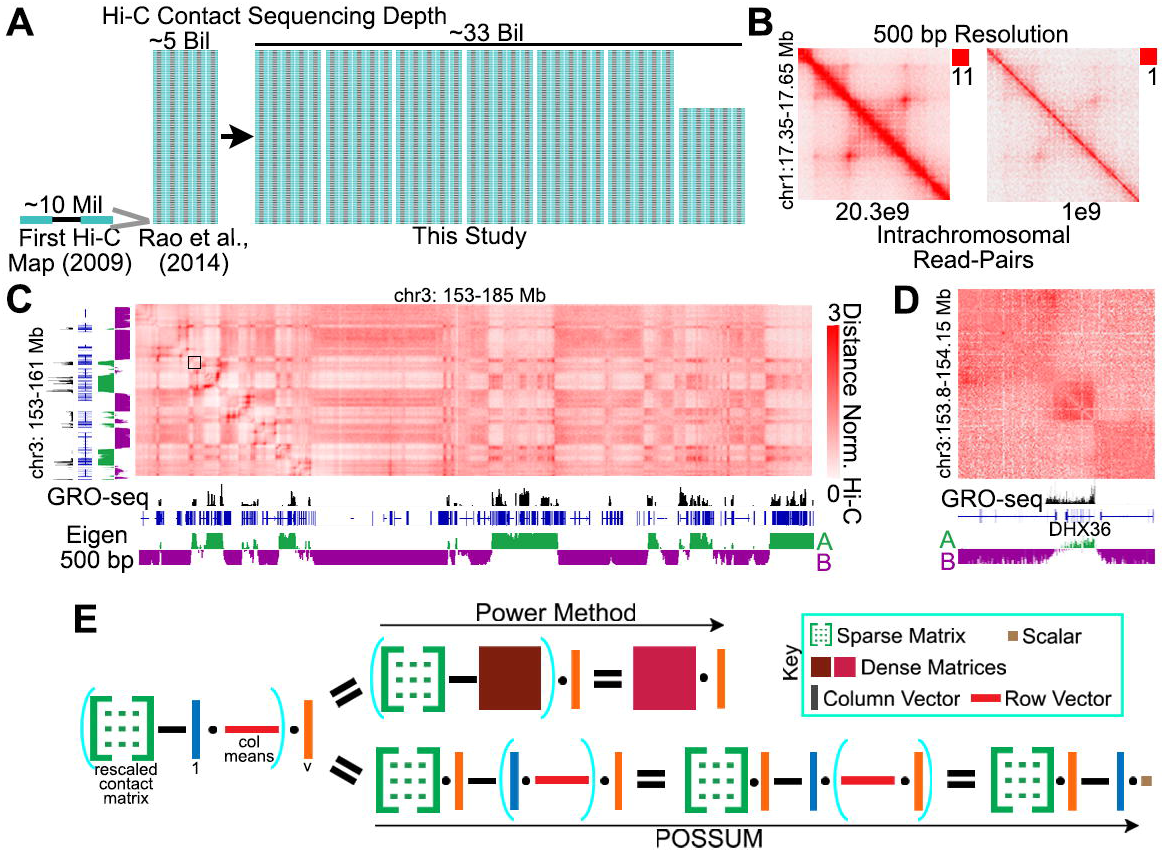
By combining ultra-deep Hi-C and POSSUMM, we generated a fine map of nuclear compartmentalization achieving 500bp resolution. A) Schematic representing the total mapped read-pairs in the current study compared to earlier published Hi-C studies. B) Example locus showing Hi-C signal in 500 bp bins in our full map with 20.3 billion intrachromosomal read-pairs (left) and when read-pairs are subsampled to 1 billion (right). C) Example of compartment interactions in a Hi-C map identified by the eigenvector in 500 bp bins (bottom track). Black track displays transcription measured by GRO-seq. Black square represents the region shown in Fig. 1D. D) Zoomed in view of a compartment domain. E) Overview of the power method and POSSUM for calculating the eigenvector. See Methods for details.

We generated contact matrices at a series of resolutions as fine as 500 bp. These matrices greatly improved the resolution of all features genome-wide, revealing many additional loops and domains (Fig. 1B). This high coverage also enhanced the long-range plaid pattern indicative of compartments (Fig. 1C, S1B), as well as the corresponding compartment domains observed along the diagonal of the map (Fig. 1D, S1C). Critically, because the number of contacts at every locus was greatly increased, with an average of 66,000 contacts incident on each kilobase of the human genome (Fig. 1C, S1B), we were able to distinguish between loci in the A compartment and loci in the B compartment with much finer resolution.

### Development of PCA of Sparse, SUper Massive Matrices (POSSUMM) and its use to create a genome-wide compartment profile with 500bp resolution

Extant methods for classifying loci into one compartment or the other typically rely on numerical linear algebra to calculate the principal eigenvector (called, in this context, “the A/B compartment eigenvector”) and the smallest eigenvalues of correlation matrices associated with the Hi-C contact matrix. At 100 kb resolution, these matrices typically have thousands of rows and columns and millions of entries, making them tractable using extant numerical algorithms, such as those implemented by Homer^8^, Juicer^9^, and Cooler^10^. However, at kilobase resolution or beyond, these matrices have hundreds of thousands of rows and hundreds of billions of entries, making them intractable using the aforementioned tools. For example, computing an eigenvector for chr1 at 500 bp resolution entails generating a matrix with 250 billion entries and performing a calculation that is projected to require >4.6 TB of RAM for >16 years (Fig. S1D).

As such, we developed a method, POSSUMM, for calculating the principal eigenvector and the smallest eigenvalues of a matrix. POSSUMM is based on the power method, which repeatedly multiplies a matrix with itself in order to calculate the principal eigenvector (Fig. 1D). However, POSSUM does not explicitly calculate all of the intermediate matrices required by the power method. Instead, it explicitly calculates only the tiny subset of intermediate values required to obtain the principal eigenvector itself, not requiring dense matrices, which makes it vastly more efficient (Fig. 1D, Fig. S1EF).

Using POSSUMM, we assigned loci to the A and B compartment at resolutions up to, and including, 500 bp (Fig. 1C). The calculation of the A/B compartment eigenvector at 500 bp resolution took only 12 minutes, and 13 GB of RAM (Fig. S1D&G). A and B compartments identified by POSSUM accurately detect the segregation of active from inactive chromatin (Fig. S1H-K).

### The median compartment interval is 12.5 kb long

It is widely thought that compartment intervals (genomic intervals that lie entirely in one compartments) are typically megabases in length and are partitioned into numerous punctate loops and loop domains^6,11–13^. To explore this phenomenon, we used our fine map of nuclear compartments to examine the frequency with which loci alternate from one compartment to the other. Nearly 99% of compartment intervals were less than 1 Mb in size, and 95% were smaller than 100 kb (Fig. 2A). The median compartment interval was only 12.5 kb, and thousands of compartment intervals were no longer than 5 kb (Fig. S1L). In comparison, the median size of CTCF loops in our map was 360 kb in length, demonstrating that compartment intervals are smaller than individual loops.

**Figure 2.**
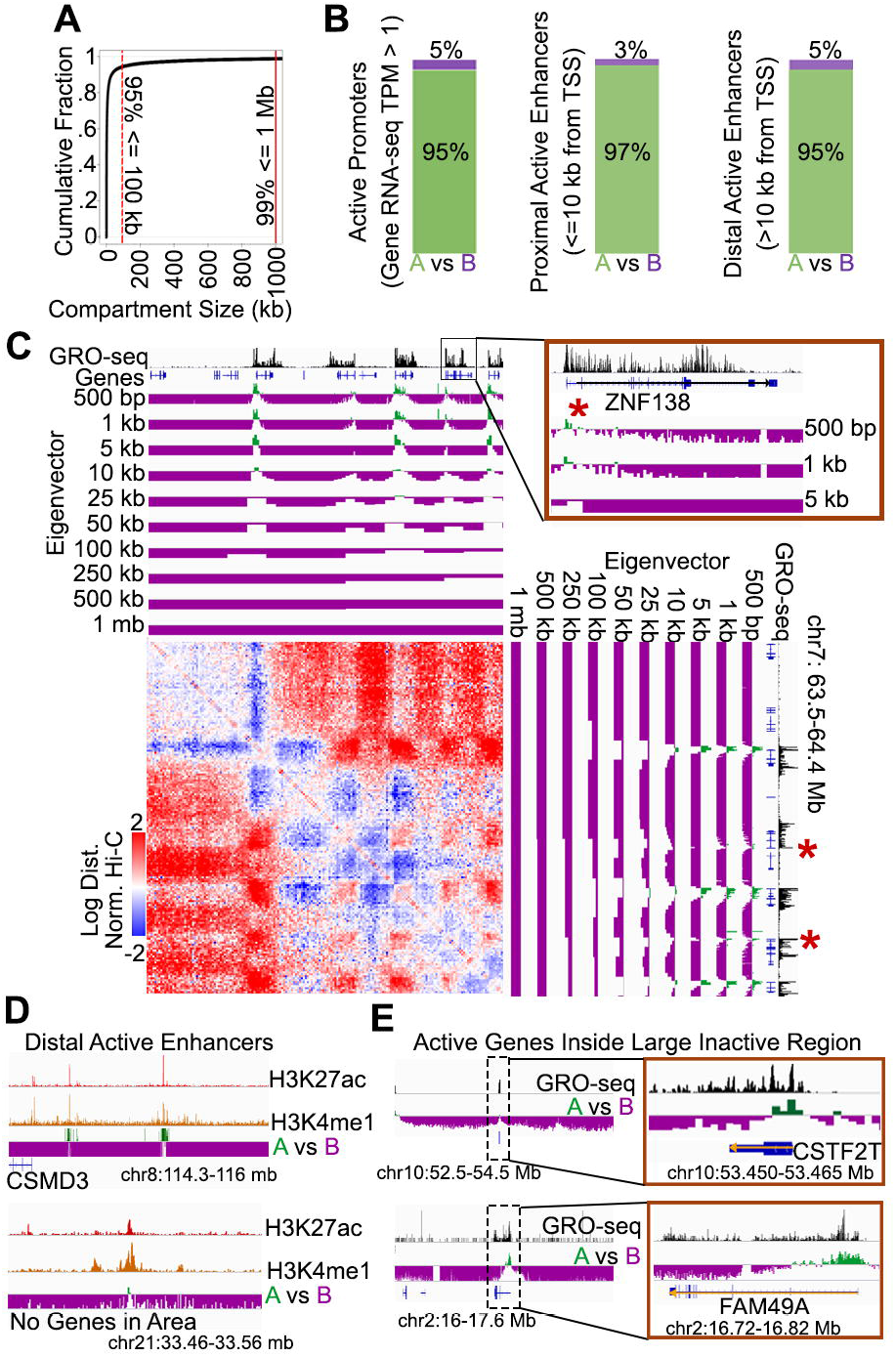
Nearly all active TSSs and Enhancers localize to kilobase-scale A compartments A) Cumulative fraction of compartment domain sizes when identified at 500 bp resolution. B) Percentage of active gene promoters, proximal enhancers, and distal enhancers assigned to A (green) or B (purple) compartment domains when identified by the 500 bp compartment eigenvector. C) Example of small compartment domains only identifiable at high-resolution (red asterisks). Log transformed and distance normalized Hi-C map is shown alongside the eigenvector tracks at various bin sizes. D) Examples an active enhancers denoted by H3K27ac and H3K4me1 signal localizing to the A compartment and surrounded by the B compartment. E) Examples an active promoters denoted by GRO-seq signal localizing to the A compartment and surrounded by the B compartment.

### Kilobase-scale compartment intervals frequently give rise to contact domains

It is well known that long compartment intervals often give rise to contact domains, i.e., genomic intervals in which all pairs of loci exhibit an enhanced frequency of contact among themselves^6,14–17^ (Fig. 1D). Such contact domains are referred to as compartment domains. We found that even short compartment intervals less than 5 kb frequently give rise to contact domains (Fig. S1M), demonstrating that intervals of chromatin in the same compartment possess the ability to form contact domains regardless of scale.

### Essentially all active promoter and enhancer elements localize in the A compartment

Next, we compared our fine map of nuclear compartments to ENCODE’s catalog of regulatory elements in GM12878 cells. We examined active promoters (defined as 500 bp near the TSS, absence of repressive marks H3K27me3 or H3K9me3, and with >= 1 RPKM gene expression in RNA-seq) and found that nearly all lie in the A compartment: out of 9,324 active promoters annotated in GM12878, only 496 (5%) were assigned to the B compartment (Fig. 2B - left). We noticed that active promoters in the B compartment had higher values in the principal eigenvector compared to the surrounding regions (Fig. S1N). Indeed, if we use a slightly more stringent threshold (assigning promoters to the B compartment only if the corresponding entry of the principal eigenvector is <−.001), we find that only 233 (2.5%) of promoters are assigned to the B compartment. Notably, when 1 Mb resolution compartment profiles are used, the number of active promoters assigned to the B compartment increases 4-fold, to ~21% (Fig. S1O). This is at least in part because the use of coarse resolutions leads to the averaging of interaction profiles from neighboring loci, such that a DNA element in the A compartment might be erroneously assigned to the B compartment if most of the flanking sequence was inactive (Fig. 2C, S2A-G).

Similarly, we found that essentially all active proximal enhancers (defined by annotation in DenDB^18^, <=10 kb from a TSS, and overlapping H3K27ac but not H3K27me3 & H3K9me3^19^) lie in the A compartment (Fig. 2B – middle). Moreover, essentially all active distal enhancers (DenDB^18^, >10 kb from a TSS, with H3K27ac, but not H3K27me3 or H3K9me3^19^) lie in the A compartment (Fig. 2B – right): out of 30,868 active distal enhancers annotated in GM12878, only 1,607 (5%) were assigned to the B compartment. Many of these distal enhancer elements represent small islands of A-compartment chromatin in a sea of inactive, B compartment chromatin (Fig. 2D). This demonstrates that individual DNA elements can escape a neighborhood that is overwhelmingly associated with one compartment in order to localize with a different compartment (Fig. 2C-E, S2H-I). When 1 Mb resolution compartment profiles are used, the number of active distal enhancers assigned to the B compartment increases 4.6-fold, to 23% (Fig. S2J). Again, this is at least in part because the use of coarse resolutions leads to the averaging of interaction profiles from neighboring loci (Fig. S2H&K).Taken together, we find that essentially all active regulatory elements, including both promoters and enhancers, lie in the A compartment, even when immediately neighboring sequences do not.

### Many genes exhibit discordant compartmentalization, with the TSS in the A compartment and the TTS in the B compartment

When exploring the fine map of nuclear compartmentalization, we noticed many genes where the TSS and TTS localize to opposite compartments (Fig. 3A., see also Fig. 1D,2C,2E). These intra-genic compartmental switches are more easily seen at large genes (Fig. 3B, S3AB). We therefore asked if gene size can affect the compartment localization of the TTS. Indeed, average profiles of compartmental status revealed that TSSs were most likely to be in the A compartment (Fig. 3C), but that the likelihood of lying in the A compartment decreases steadily as one examines increasingly distal portions of the gene body, such that the TTSs of large genes are more likely to localize to the B compartment (Fig. 3C&D, S3C). This was especially evident if we consider very large genes (Fig. S3D), where the TSS was overwhelmingly in the A compartment, but the TTS was usually in the B compartment.

**Figure 3.**
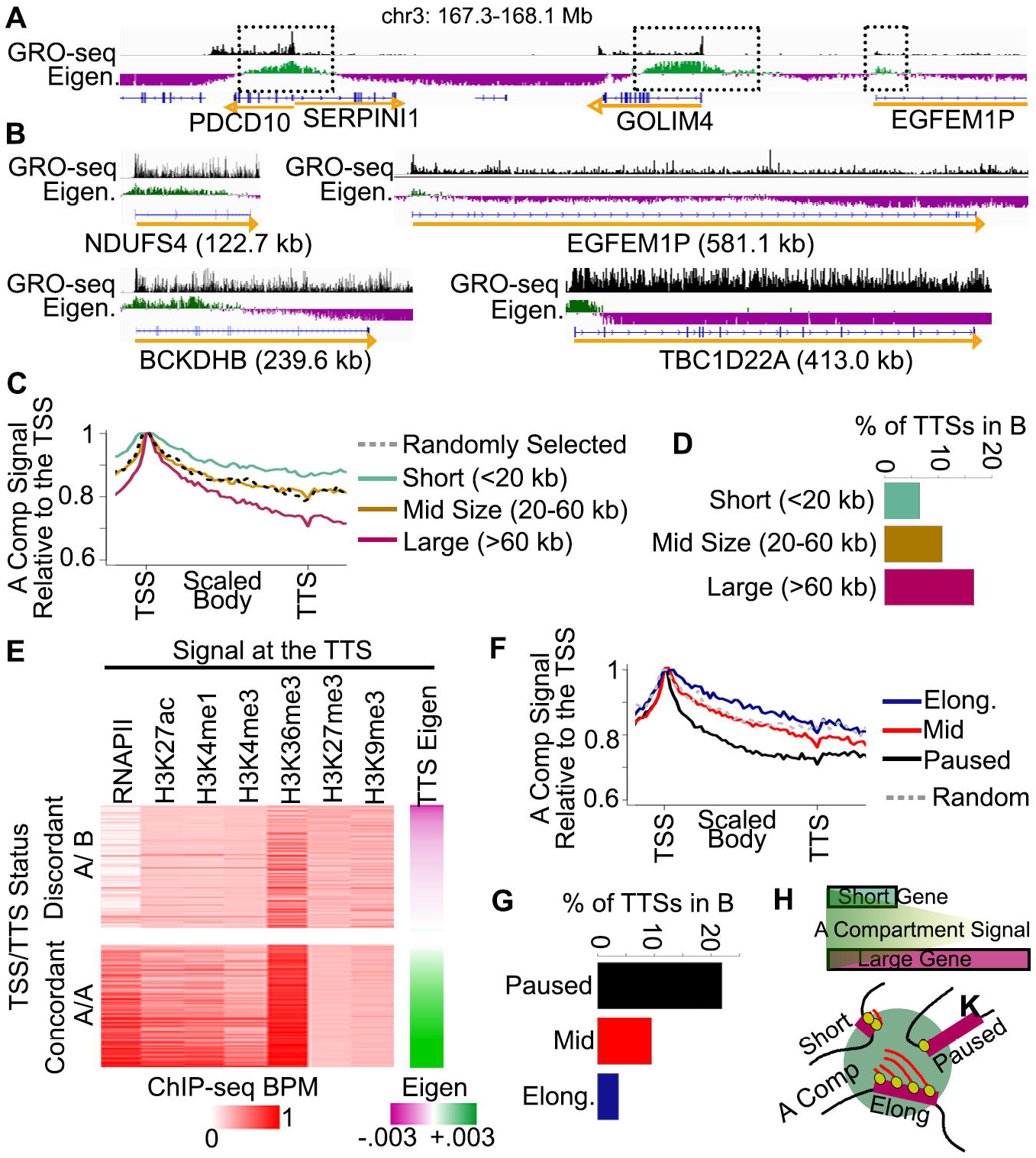
Many genes exhibit discordant compartmentalization. A-B) Examples of genes of various sizes where the TSS is in the A compartment while the TTS is in the B compartment. GRO-seq signal is shown as an indicator of the gene’s transcription status. C) Scaled average profiles of the A compartment signal (positive eigenvector) relative to the TSS for short (blue), mid-sized (gold), large (pink), and randomly selected (black) genes. D) Percentage of TTSs that localize to the B compartment for genes of various sizes (left). E) ChIP-seq signal at the TTS of discordant A/B genes vs. concordant A/A genes. Genes are sorted by the TTS compartmental signal. F) Scaled average profiles of the A compartment signal (positive eigenvector) relative to the TSS for elongating (blue), mid (red), paused (black), or randomly selected (grey) genes. G) Percentage of TTs that localize to the B compartment for paused, mid, or elongating genes. H)Top: Simple diagram of A compartment signal relative to gene size. Bottom: Diagram of TSS and TTS localization to the A compartment depending on gene size and elongation status.

We next asked if genes with discordant compartmentalization (i.e., the TSS was in compartment A, but the TTS was in compartment B) could be explained by different chromatin marks at the TSS vs. TTS. We examined chromatin marks at the TTS in active genes larger than 20 kb and found that diminished levels of active marks at the TTS, specifically RNAPII, H3K4me1, and H3K36me3, were correlated with presence of discordant compartmentalization (Fig. 3E, Fig. S3E). Notably, although repressive chromatin marks are frequently seen at loci in the B compartment, genes with discordant compartmentalization typically lacked such marks at the TTS (Fig. 3E, S3E). We also found that chromatin marks at the TSS were not predictive of whether the gene exhibited discordant compartmentalization (Fig. S3E&F).

Finally, we sought to determine if discordant compartmentalization was associated with transcriptional pausing as measured by GRO-Seq. We found that elongating genes longer than 20 kb were more likely to exhibit concordant compartmentalization (Fig. 3F), whereas paused genes were more likely to exhibit discordant compartmentalization (Fig. 3G).

Taken together, these data support a model where an active TSS localizes to the A compartment but brings with it only a small portion of the gene body, depending on the elongation status (Fig. 3H).

### Loop extrusion forms diffuse loops, whereas compartmentalization forms punctate loops

We examined loops in our Hi-C dataset. Using SIP^20^ and HiCCUPS^2^, we identified 32,970 loops. Ninety-one percent of these loops contained a CTCF-bound motif at both anchors, with a strong preference for the convergent orientation (Fig. S4A).

Interestingly, when we examined loops at 1 kb resolution, we noticed that the signal is diffuse (Fig. 4A, S4B), indicative of frequent contacts proximal to the CTCF binding sites (Fig. 4B). The elevated contact frequency decays as the distance from the corresponding anchors increases (Fig. 4C, rainbow) (a loss of signal of c.a. −6% from one bin to the next; i.e. −6%/kb compounding). Curiously, this decay rate is much slower than the decay rate reflected by the diagonal of the Hi-C map (Fig. 4C, S4C – expected) (c.a. −28%/kb), which is thought to reflect the properties of the chromatin polymer. The decay was unchanged as a function of loop size or sequencing depth (Fig. S4DE).

**Figure 4.**
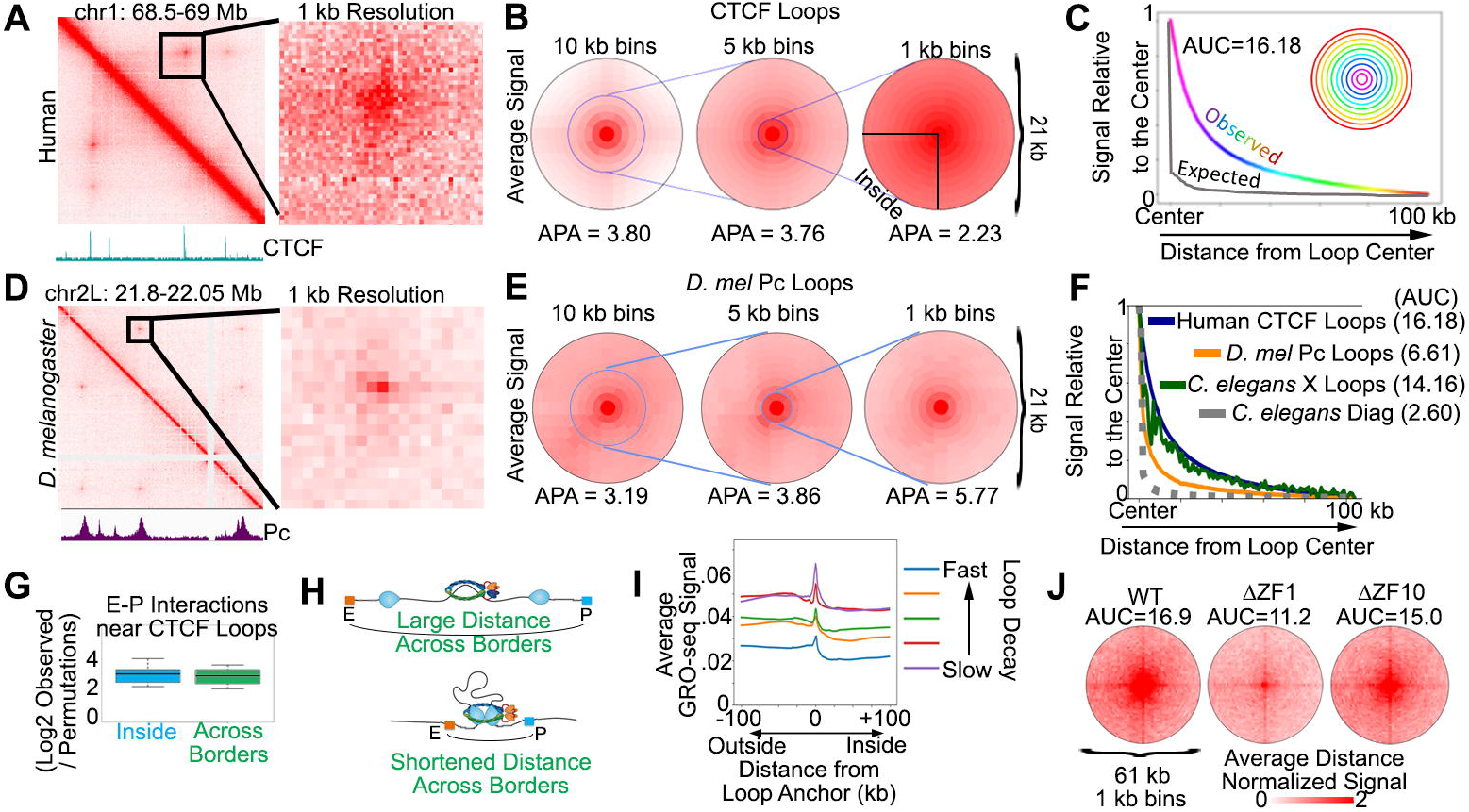
CTCF loop-decay enhances proximal interactions and is dependent on RNA-binding domains. A) Example of broad signal enrichment near CTCF loops when binned at 1 kb. B) Average signal at CTCF loops when binned at 10, 5, or 1 kb, centered on convergent CTCF anchors. C) Average Hi-C signal in 1 kb bins at each radial distance away from the CTCF loop anchors (rainbow). Average signal of the diagonal decay is shown for reference (grey) to estimate interactions due to polymeric distance. AUC=area under the curve. D) Example of punctate signal enrichment at Pc loops in *D. melanogaster* when binned at 1 kb. E) Average signal at *D. melanogaster* Pc loops when binned at 10, 5, or 1 kb. F) Average Hi-C signal in 1 kb bins at each radial distance away from human CTCF loop anchors (blue) vs. D. melanogaster Pc loops (orange), and *C. elegans* X-chromosome loops (green). Average signal at the *C. elegans* Hi-C diagonal is shown for reference (grey). AUC=area under the curve. G) Enrichment of Fit-Hi-C enhancer-promoter interactions within 100 kb of loops inside the loop (blue) or crossing over loop boundaries (green). Values are shown as enrichment vs random regions of equal size and number as loops. H) Diagram of how CTCF loops can shorten distances between enhancers (orange) and promoters (blue) even when both are located outside of the loop. I) Average GRO-seq signal at CTCF loop anchors and neighboring loci for loops divided into 5 distinct decay categories. J) Average Hi-C signal in WT (left), ΔZF1 (middle), or ΔZF10 (right) CTCF mutants at CTCF loops. AUC=area under the curve

We wondered whether this slow decay in contact frequency was seen for loops in other species. We therefore examined hundreds of loops observed in a published high-resolution Hi-C map from *Drosophila melanogaster* Kc167 cells at 1 kb resolution^14,21^ (Fig. 4D&E). Interestingly, the loops in *Drosophila* decayed at a rate (c.a. −20%/kb) that matched the diagonal of the *Drosophila* Hi-C map (c.a. - 23%/kb) and was much faster than the rate seen for human CTCF-mediated loops (Fig. 4F). This suggests that CTCF loops create interactions between sequences bound by CTCF, as well as interactions between CTCF bound and adjacent sequences. However, in *Drosophila*, Pc loops only create interactions directly between the Pc bound sequences.

Finally, we examined loops previously identified in *C. elegans*^20,22,23^. The loop decay was slower (c.a. −11%/kb) than the decay seen at the diagonal (c.a. −24%/kb) (Fig. 4F, green vs. grey), and was more similar to the rate of decay seen for human CTCF-mediated loops than the one observed for *D. melanogaster* loops (Fig. 4F, Fig. S4I).

It was notable that the type of decay observed (fast or slow) matched the putative mechanism by which the loops formed. CTCF-mediated loops in human are bound by, and dependent on, the SMC complex cohesin (Fig. S4H), and form by cohesin-mediated extrusion^24–27^. Similarly, the loops in *C. elegans* are bound by the SMC complex condensin and we previously suggested that they are formed by condensin-mediated loop extrusion^20,22,23^. Indeed, the interactions between loop-adjacent sequences are in further support of loop formation by extrusion in *C. elegans*. By contrast, Drosophila loops are much less likely to be bound by CTCF, cohesin, condensin, or other extrusion-associated proteins^14^. Instead, they are bound by the Polycomb complex, Pc, and may form by means other than extrusion^28–30^.

These findings suggest that the mechanism of loop formation influences whether loops will be punctate or diffuse, with extrusion-mediated loops forming diffuse peaks and compartmentalization-mediated loops forming more punctate features.

### Diffuse loops enhance the contact frequency of nearby promoter-enhancer interactions

Using Fit-Hi-C^31^, we called promoter-enhancer interactions at 1 kb resolution on human chr1. We examined those interactions where both the promoter and enhancer lie within 100 kb of a loop anchor. In some cases, these interactions lie completely inside the loop, but in others they cross the loop anchor. Both cases exhibited strongly enriched contact frequency as compared to enhancer-promoter interactions that are unrelated to CTCF loops, i.e., near permutated random sites (Fig. 4G). These data suggest that CTCF loops enhance the contact frequency of promoter-enhancer interactions, even when both elements lie outside the loop (Fig. 4H). By contrast, in *Drosophila*, Fit-Hi-C interactions between promoters and enhancers tend to be much shorter (Fig. S4J).

### Deletion of CTCF’s RNA binding domains leads to more punctate loops

Interestingly, we observed some variability in the decay rate for different loops (Fig. S4K). This decay did not correlate strongly with either CTCF motif strength, CTCF ChIP-seq peak strength, or Rad21 ChIP-seq peak strength (Fig. S4L-O). Instead, we found that CTCF-mediated loops exhibiting slower decay are associated with higher levels of transcription (Fig. 4I) and chromatin accessibility (Fig. S4P) near the loop anchors. This suggests that nearby transcriptional activity could impact how CTCF interacts with the nearby sequences and / or with the loop extrusion process.

The CTCF protein contains 11 zinc finger domains. Recently, it was shown that ZF1 and ZF10 bind to RNA, and that deletion of these two domains causes weakening of loops throughout the genome^32^. We performed aggregate peak analysis on the published Hi-C in ZF1 and ZF10 mutants^32^ using “bullseye” plots in order to explore the effect of these deletions on loop decay. Interestingly, we found that loops appeared more punctate in both CTCF RNA binding mutants (Fig. 4J). This effect was especially pronounced in the ZF1 mutant.

Taken together, these findings are consistent with a model where CTCF’s RNA-binding domains and the presence of bound RNAs results in more diffuse contacts between loop anchors, and thus to enriched contacts among regulatory elements near the loop.

## Discussion

By generating a Hi-C map with extraordinary sequencing depth (33 billion PE, or 9.9 terabases of uniquely mapped sequence), we create the first fine-map of nuclear compartmentalization.

Our findings demonstrate that compartment intervals and compartment domains can be far smaller than previously appreciated. This contrasts with the common hierarchical model of chromosome organization in which compartments are partitioned into TADs and loops^6,11–13^. In fact, our results indicate that compartment intervals can be so small that active DNA elements will localize with the A compartment even when surrounded by inactive chromatin localizing in the B compartment (Fig. 5).

**Figure 5.**
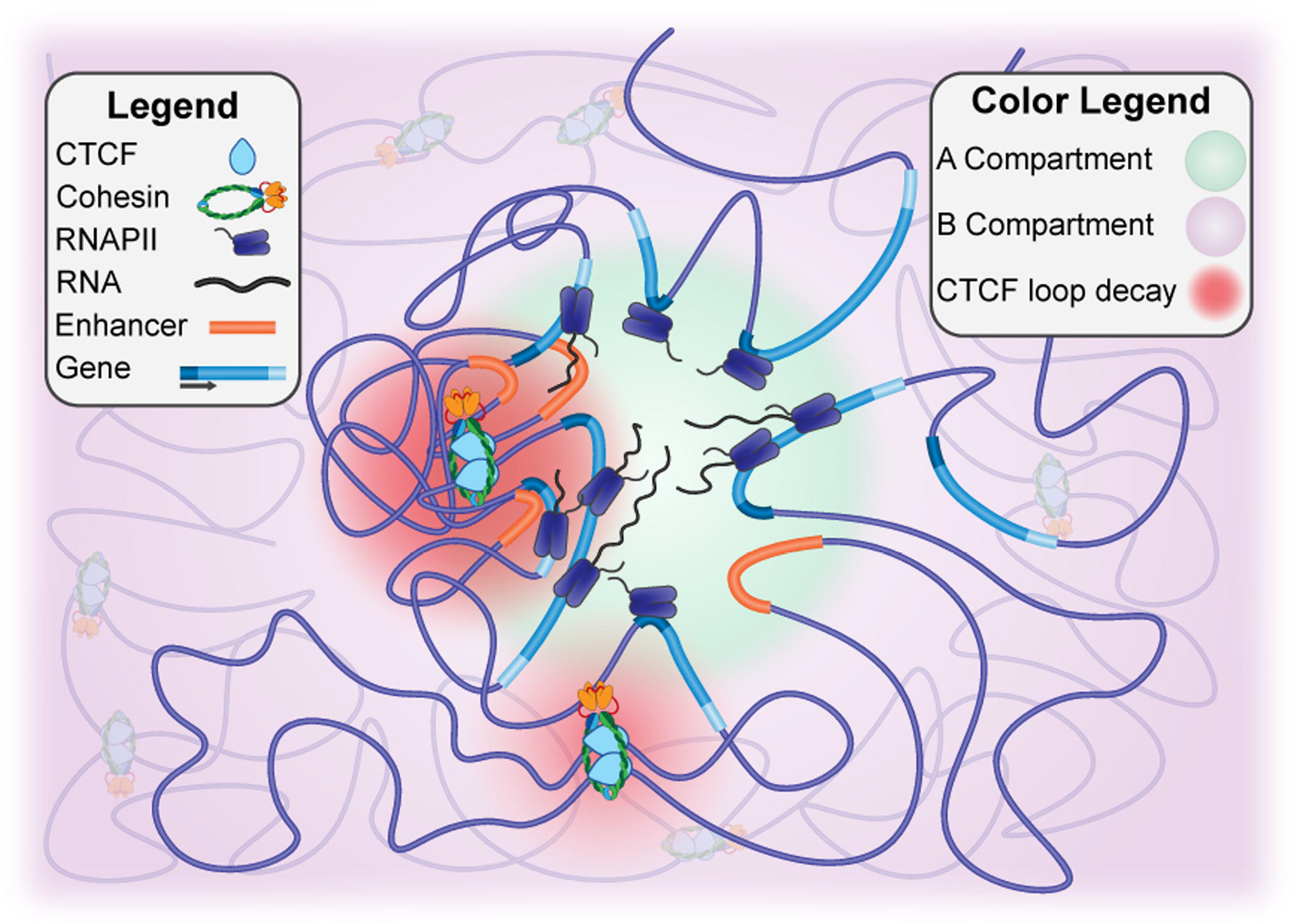
Sub-genic compartmentalization and diffuse CTCF looping organize the human genome. Diagram depicting localization of active enhancers and TSSs to the A compartment, while TTSs are oriented to the B compartment dependent on transcription elongation status. This sub-genic and precise enhancer compartmentalization combines with diffuse CTCF loops to mediate genome organization.

Strikingly, we find that essentially all distal enhancer elements lie in the A compartment. This contrasts with earlier work, using coarse-resolution maps of compartmentalization, which only report general enrichment of active distal enhancers in the A compartment, rather than as an absolute characteristic of active enhancers^33,34^. Similarly, many previous studies have reported a coarse enrichment of active genes in the A compartment^6^, yet we find that essentially all active promoters lie in the A compartment.

We also observe that the likelihood that a locus lies inside the A compartment declines as one moves away from the promoter, along the gene body. Interestingly, we observe numerous genes with discordant compartmentalization, where the TSS and TTS tend to be in different compartments. This observation suggests that opposing compartments need not correspond to widely separated locations within the nucleus. For instance, recent work indicates that compartments could be phase-separated droplets^35^, suggesting that the TSS and TTS of a gene with discordant compartmentalization might be physically proximal within the nucleus, in neighboring A and B droplets (Fig. 5).

The finding that active promoters – specifically, active TSSs – are overwhelmingly localized in the A compartments; that TTS compartment status correlates with RNAPII levels at the TTS; and that genes with discordant compartmentalization tend to be transcriptionally paused is consistent with a model in which RNAPII drives localization to the A compartment. Although a recent RNAPII degradation study showed little effect on genome organization, these experiments did not achieve the sequencing depth required to perform fine mapping of nuclear compartmentalization, nor to resolve phenomena such as genes with discordant compartmentalization^36^. Alternatively, other components of the transcription complex that travel along the gene body during transcription elongation may be responsible for mediating interactions that assign sequences to the A compartment. In future studies, it will be of great interest to examine how RNAPII and other components of the transcription complex impact genome organization at the TSS and TTS separately.

We note that our data represent averages within the cellular population, and it is unclear where each component lies during the transcriptional process itself. In the future, fine mapping of nuclear compartments in single cells will be needed to decipher these dynamics. Moreover, we note that our study did not attempt to study subcompartments or models with >=3 distinct compartment states^2,37^, which will be an important topic for future analyses.

Our ultra-deep Hi-C map also helped identify interesting properties of chromatin loops. In particular, we observe that CTCF-mediated loops are highly diffuse, more so than would be predicted based on polymer behavior alone (Fig. 5). Interestingly, this diffusivity is observed for loops that form by extrusion, such as loops in human^2,24–27^ and *C. elegans*^20,22,23^, but is not observed for loops that are believed to form by compartmentalization, such as the numerous Pc-associated loops observed in *Drosophila*^14,21,29,30^. Intriguingly, variations in diffusivity between different loops could explain differences in domains signal (See Supplemental Discussion, Fig. S5).

*In vitro* studies have found that large chromatin complexes can impede looping factors^38,39^, and cohesin was shown to build up near transcriptionally active regions^40^. Yet studies have also reported independence of CTCF loops and transcription^36,41,42^, bringing the relationship between transcription and CTCF looping in question. Recently, it was shown that CTCF RNA-binding domains, ZF1 and ZF10, are important for looping^32^. Our finding that loop-decay is altered in CTCF RNA-binding mutants supports the argument that transcription can impact fine-scale chromatin organization in mammals, as does the correlation between TTS compartmental domains and elongation status.

Our POSSUMM method, a novel numerical linear algebra algorithm for calculating principal eigenvectors, is now available as part of the Juicer pipeline for Hi-C analysis. Our power analyses suggest that fine mapping of nuclear compartments at sub-kilobase resolution becomes possible for maps containing 7 billion contacts or more (See Supplemental Discussion, Fig. S6&S7). As sequencing costs continue to decline, we expect that fine mapping of nuclear compartments will become increasingly common.

## Methods

### Library Preparation, Initial Processing, and Quality Metrics

Hi-C libraries were prepared according to the published in-situ method^2^. The full map represents a mixture of libraries prepared by digestion of various 4-cutter restriction enzymes, MboI, MseI, and NlaIII. Reads were aligned to the hg19 genome, processed, Knight-Ruiz (KR) normalized using Juicer^9^. Subsampled Hi-C maps were created by uniform random selection of read-pairs from the 33.3 billion Hi-C dataset. We provide a script for subsampling Hi-C data at https://github.com/JRowleyLab/HiCSampler.

### Compartment Analysis

Compartments were identified using the A/B eigenvector of the Hi-C matrix using POSSUM. POSSUMM can be downloaded from: https://github.com/aidenlab/EigenVector and is also now implemented in the ENCODE version of the Juicer pipeline: https://github.com/ENCODE-DCC/hic-pipeline.

## Introduction to PCA of Sparse, SUper Massive Matrices (POSSUM)

We note that the so-called “A/B compartment eigenvector” is simply the eigenvector of A corresponding to its largest eigenvalue, where X is given by the Hi-C contact matrix. This is equivalent to the first principal component in Principal Component Analysis. We note that in our case, X is a large, sparse matrix, containing millions of rows, millions of columns, and tens of billions of nonzero entries (dubbed a “Sparse, SUper Massive Matrix”).

Suppose we seek to calculate the largest eigenpairs, *λ*_*i*_, *v*_*i*_ of A in this case. Although X is sparse, we note that both Y and A are dense matrices. Unfortunately, storing dense matrices with millions of rows and columns in memory is impossible. Hence we cannot use any method for calculating the eigenvectors of A that would require us to explicitly calculate either Y or A. Similarly, traditional sparse matrix methods for eigendecomposition are not usable here, again because A - the correlation matrix we hope to analyze - is a dense matrix.

Therefore, in order to calculate eigenvectors for A, we began by implementing a method that makes it possible to calculate the matrix-vector product Av (where v is an arbitrary vector) using a sparse representation of X, i.e., without explicitly computing either A or Y. See POSSUMM details below for a more complete description.

Next, we note that there are many methods for calculating eigenvectors in which the input matrix only appears via a matrix-vector product. These include the Power method, the Lanczos method, and their many variants^43^. Thus, in principle, any of these methods - for which there are many implementations in Fortran, C, C++, Matlab, and R - can be combined with the sparse Av product calculation described above in order to calculate eigenpairs of A. In practice, methods combining these two approaches are not available.

To the best of our knowledge, the sole exception is a method in the R package irlba, which was released while this study was being performed. The details of this method are unpublished, but the method itself is available at https://cran.r-project.org/web/packages/irlba/index.html. However, irlba cannot handle cases where X has more than roughly two billion nonzero entries, which is exceeded in the present case. It also does not enable parallelization, which limits performance in highly demanding settings.

POSSUMM combines sparse Av product calculation with the power method, is extremely memory-efficient, and enables parallelization via multi-threading.

## POSSUMM Details

To identify compartments from sparse Hi-C matrices, we began by excluding all rows and columns with 0 variance. Let *X* be a matrix with column vectors *X*^(1)^,…, *X*^(*n*)^. Let *Y*^(*i*)^ = (*X*^(*i*)^ - *c*_*i*_)/*σ*_*i*_ 1 ≤ *i* ≤ *n*, where *c*_*i*_ is the mean of *X*_*i*_ and *σ*_*i*_ is its standard deviation. Let *Y*= (*Y*^(*i*)^, …, *Y*^(*n*)^) be an n × n matrix with column vectors. The correlation matrix of *X* is *A*= *Y*^*T*^*Y* where *Y*^*T*^ is transposed *Y*. Since A is symmetric and positive semi-definite it has n real eigenvalues *λ*_1_ ≥ *λ*_2_ ≥ … ≥ *λ*_*n*_ ≥ 0 and *n* eigenvectors. *v*_1_,…, *v*_*n*_ where *Av*_*i*_ - *λ*_*i*_*v*_*i*_.

These eigenvectors are a basis of *R*^*n*^ (i.e., a set of vectors which are independent and span the space) if *λ*_*i*_ ≠ *λ*_*j*_ and *v*_*i*_ ⊥ *v*_*i*_ (i.e., 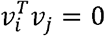) . To compute *v*_1_ using the power method (a.k.a power iterations), suppose that *λ*_1_ > *λ*_2_and let *x*_0_ be any nonzero vector in *R*^*n*^, we define the recursive relation:
*x*_*k*+1_ = *Ax*_*k*_= *A*^*k*+1^*x*_0_. We can represent *x*_0_ as *x*_0_ = *a*_1_*v*_1_ + … +*a*_*n*_*v*_*n*_ and therefore 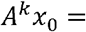 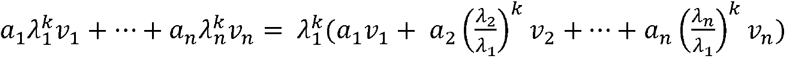. Once we have estimates of the eigenvector and the two largest eigenvalues, we can estimate the error given that ||*v* − *v*_1_|| ≤ 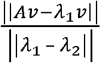. To find an estimate of *λ*_2_ we know that *v*_2_ ⊥ *v*_1_ and ||*v*_1_|| = 1. Let *x*_0_be any vector and let *x*_*k*+1_ =*A*(*k*_*x*_ − *c*_*k*_*v*_1_) where 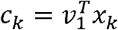 (and then (*x*_*k*_ − *c*_*k*_*v*_1_) ⊥ *v*_1_). If 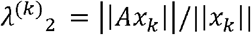 the using the same argument as before 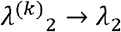 as *k* → ∞. This is true even if *λ*_*2*_ ≈ *λ*_*3*_ (*x*_*k*_ may not converge to *v*_2_, but *λ*_2_ wil converge to *λ*_2_). In this way we have an estimate of *λ*_1_ and *λ*_2_ and may estimate the error in *v*. Since *A*= *Y*^*T*^*Y*, *Ax* =*Y*^*T*^(*Yx*) = ((*Yx*)^*T*^*Y*)^*T*^, we do not need to compute A (which has the complexity of *O*(*n*^3^)). We used two matrix vector products at every iteration (which have the complexity of the number of nonzero elements in Y which is at most *O*(*n*)). Moreover, if *X* is large a naïve multiplication of a vector by a matrix can still take a long time and storing *Y* may require a large amount of memory. For example, to store human chr1 at 1 kb resolution (where *n* ≈ 250000) 500 GB of RAM would be required just to store *Y*. With sparse implementation we recall that *Y*= (*Y*^(*i*)^,… , *Y*^(*n*)^) where 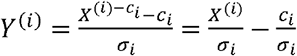. While 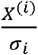 is sparse, 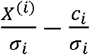 is not. In lieu of explicit computation, let 1 = (1,1, … ,1)^*T*^ then 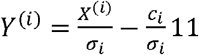 and then *Y* = *XS* − 1 · 1 · *r*^*T*^ where 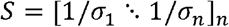 and 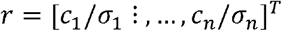 and then *Yx* - (*X* · *S*)*x* − 1 ·*r*^*T*^ ·*x*. Let *Z*= *X* · *S*. Since 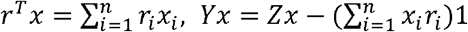. Since Z is as sparse as X we can do everything with sparse matrices as 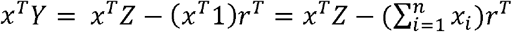. Projected time and memory usage were calculated by fitting a power decay curve, R^2^ of fit = 0.95 for time, and R^2^ of fit = 0.98 for memory usage.

After compartment calling, chromatin marks were profiled at features that overlap A or B compartments by overlapping with ChIP-seq peaks and by using average signal profiles created by pyBigWig from the deepTools package^44^. ChIP-seq peaks and bigwig files were obtained from the ENCODE Roadmap Epigenomics project^45^. We filtered promoters with bivalent marks as active genes that had 2-fold higher H3K27me3 or H3K9me3 signal compared to the average at promoters. Contiguous compartment domain sizes were calculated by requiring at least two consecutive bins to have the same sign in the eigenvector. To create profiles of A compartmental status along genes, we assigned genes to elongating, mid, and paused. Elongation status was determined by RPKM GRO-seq signal within 250 bp of the TSS compared to the gene body, excluding 500 bp from the TSS. Differences between Promoter – Gene Body GRO-seq signal were ranked and placed into three equal categories considering only genes >= 20 kb in size.

### Loop Analysis

Loops were identified by HiCCUPS^2^ or SIP^20^ at multiple resolutions. For HiCCUPS, we used parameters – m 2000 –r 500,1000,5000,10000 –f .05,.05.05.05. For SIP we used an FDR 0.05 at each resolution with the parameters for resolutions of 500 bp; -d 15 –g 3.0; 1 kb: -d 17 –g 2.5; 5 kb: -d 6 –g 1.5; and 10 kb: -d 5 –g 1.3. Loops called by both methods were combined by placing all loops into 10 kb bins, and if HiCCUPS and SIP called the same loop within the 10 kb bin, then only one instance of this loop was kept. Loops in subsampled maps were overlapped with loops called in the full 20.3 billion map if the loop was within +/− 25 kb of each other. Overlap of loops with CTCF was done using a published list of CTCF ChIP-seq peaks and motifs^2^. Central 1 kb bins were assigned to those where we could unambiguously assign a CTCF ChIP-seq peak to a unique bin at motifs in convergent orientation. Only loops with unambiguous CTCF assignment were used in decay analysis. Bullseye plots were created using SIPMeta^20^ and the decay was calculated as the average at each Manhattan distance (ring) moving away from the central bin. These values were plotted as a ratio to the central bin’s signal. The central bin of loops called at AUC values were computed using Simpson’s rule. Loops were placed into five equally sized categories (quintiles) based on AUC values. AUC values between WT, ΔZF1, and ΔZF10 were normalized by the diagonal to account for differences in the expected decay. The decay percentage rate of change listed in the main text was calculated by averaging the number of kb between each 10% loss of signal.

Fit-Hi-C^31^ interactions were identified in 1 kb bin-pairs with an FDR 0.05. 3D loop models were created with Pastis^46^ using the raw Hi-C matrix. Models were visualized in ChimeraX^47^.

### Comparison with Other Datasets

Hi-C read-pairs from CTCF ΔZF1, ΔZF1, and wild-type were downloaded from GSE125595^32^ and processed with juicer to the mm10 genome. Hi-C maps from the *D. melanogaster* dm6 genome and the *C. elegans* ce10 genome were obtained from our previously published work^20,21^. Hi-C maps used in our metric comparison are listed in Tables S2 and S3.

Enhancers were downloaded from DENdb^18^ and active enhancers were defined as those that overlap with H3K27ac ChIP-seq peaks in GM12878. Histone modification ChIP-seq data was obtained from the ENCODE reference epigenome series (ENCSR977QPF) and RNAPII ChIP-seq peaks were combined from RNAPII, RNAPIISer2ph, and RNAPIISer5ph (ENCSR447YYN and ENCSR000DZK)^19,48^, with overlapping peaks merged into a single peak. GRO-seq data from GM12878 was downloaded from GSM1480326^49^, and chromHMM states for GM12878 were downloaded from the Roadmap Epigenomics Project^45^.

## Data and Code Availability

Hi-C data can be downloaded from ENCODE Accession: ENCSXXXXX. Our programs for subsampling, noise estimation, and eigenvector calculation on sparse matrices can be downloaded from https://github.com/JRowleyLab/HiCSampler, https://github.com/JRowleyLab/HiCNoiseMeasurer, and https://github.com/aidenlab/EigenVector. These are open source and include source code as well as implementations in python and C++.

## Acknowledgements

We acknowledge additional members of the ENCODE consortiums Nuclear Architecture Working Group for thought-provoking discussions. Research reported in this publication was supported by a Cornelia de Lange Syndrome Foundation grant and the National Institutes of Health (NIH) under Award Numbers T32-GM067553, R35-GM139408, R35-GM128645, R01-MH115957, and U24~HG009446. E.L.A. was supported by the Welch Foundation (Q-1866), a McNair Medical Institute Scholar Award, an NIH Encyclopedia of DNA Elements Mapping Center Award (UM1HG009375), a US-Israel Binational Science Foundation Award (2019276), the Behavioral Plasticity Research Institute (NSF DBI-2021795), NSF Physics Frontiers Center Award (NSF PHY-2019745), and an NIH CEGS (RM1HG011016-01A1). M.J.R was supported by the NIH National Institute of General Medical Sciences (NIGMS) Pathway to Independence award R00-GM127671. The content is solely the responsibility of the authors and does not necessarily represent the official views of the NIH.

## Author Contributions

H.G prepared Hi-C libraries for sequencing with samples prepared by S.K., K.M., M.E.T, and J.S.S. H.G., H.H., Y.E., A. Krishna, A. Kalluchi, M.P., S.S.P.R., O.D, D.U., M.H.N., and E.D. contributed ideas and in performing various quality metrics. M.J. and G.C. created 3D loop models. M.O. created POSSUM. D.H.P., V.G.C, W.S.N, E.L.A., and M.J.R. supervised the work and wrote the manuscript. All other analyses were performed by M.J.R.

## Ethics Declarations

We declare that the authors have no competing interests in this work.

