## Supplemental Material for "Fine-mapping of nuclear compartments using ultra-deep Hi-C shows that active promoter and enhancer elements localize in the active A compartment even when adjacent sequences do not"

**Table of Contents**

Supplemental Discussion

Supplemental Methods

Supplemental Figures S1 to S7

Supplemental Tables S1 to S4

Supplemental References

### Supplemental Discussion

#### Loop-Decay Combined with Polymer Decay can Explain CTCF Loop Domains

*Drosophila* Pc loops do not create strong domains (i.e., strong signal inside of loops)[1-4], contrasting with CTCF loops in human cells. These differences in domains can be seen by 3D modeling of individual loops (Fig. S5A). Therefore, we investigated the potential impacts of loop decay on loop domains. Some CTCF loops create strong contact domains while others have weaker domains (Fig. S5B), and in examining individual loops we noticed that slowly decaying loops correlate with stronger domain-like features (Fig. S5C). We also found that domain calls tend to skew towards the smaller loops (Fig. S5D-F). By plotting the average signal inside loops starting near the diagonal and outward, we identified an inflection point where the signal increases as it approaches the interacting CTCF anchors (Fig. S5G). This inflection point is likely where the diagonal decay meets the radial loop decay (Fig. S5H). Indeed, Hi-C distance normalization leaves only the loop decay (Fig. S5IJ). We propose that interactions between functional elements decay with linear distance, and that CTCF loops have evolved to reassert an interaction decay at distal loop anchors. Therefore, linearly proximal functional elements which are also proximal to CTCF loop anchors will be subject to a combination of these two decaying interaction signals (Fig. S5H).

#### Hi-C Sequencing Depth Guidelines

Because this Hi-C sequencing effort represents 90x higher coverage than the average published map (Fig. S1A), we next tested the effects of coverage on analysis, with the aim of establishing sequencing depth guidelines for Hi-C data. We began by randomly subsampling read-pairs to create Hi-C maps with various sequencing depths. While the slope of diagonal decay (i.e., the average signal at each distance) was unchanged (Fig. S6A), the information content in bin-pairs was dependent on sequencing depth (Fig. S6B-E). For example, when examining all bin-pairs within 1 Mb of each other, 3 billion intra-chromosomal read-pairs are required to achieve a frequency at which 90% of the bin-pairs have at least 1 read (Fig. S6B). We also found that Hi-C heatmaps from samples with low read coverage result in bigger signal differences between neighboring bin-pairs, creating noisier maps (Fig. S6F-H). Notably, by these two metrics, published Hi-C maps with less than 3 billion intra-chromosomal read-pairs have low information content and high noise levels when placed in bins smaller than 5 kb (Fig. S6C,E,H).

We next tested how sequencing depth affects feature identification. Sequencing depth dramatically influences long-range compartment interactions (Fig. S7A) and maps with <7 billion intra-chromosomal read-pairs failed to identify compartments at 500 bp resolution (Fig. S7B). By testing other bin sizes, we surprisingly found that only 100 million intra-chromosomal read-pairs are needed to identify A/B compartments at 25 kb resolution with 95% consistency (Fig. S7C). Therefore, compartment identification can be done at higher resolutions than the usual 1 Mb or 100 kb bins even with Hi-C data sequenced at low depth. Based on these data we suggest the minimum intra-chromosomal read-pairs is 7 billion for 500 bp, 2 billion for 5 kb, and 250 million for 10 kb resolved compartments (Fig. S7C).

Upon examining CTCF loops in subsampled maps, we found that low sequencing depth resulted in lower signal-to-background ratios measurable by aggregate peak analysis (APA) (Fig. S7D-F). As such, sequencing depth correlates with the number of identifiable loops (Fig. S7G). For example, we estimate that *in situ* Hi-C maps with <=500 million intra-chromosomal read-pairs, i.e. 90% of published maps (Table S2), may be missing more than 57% of loops simply due to low coverage (Fig. S7H). However, we found that sequencing depth does not dramatically impact the potential false-positive rate (Fig. S7I). We investigated why some loops are missed with lower sequencing depths, finding that convergent motifs are prominent on loops called at all depths (Fig. S7J). Instead, sequencing depth correlates with the ability to call loops at longer distances (Fig. S7K). Most published Hi-C datasets have less than 500 million intra-chromosomal contacts, suggesting that these data were unable to detect the majority of CTCF loops larger than 1 Mb. Based on the plateau of the distance bias (Fig. S7K), we estimate that only 5% of CTCF loops are greater than 3.4 Mb in size (Fig. S7L).

Our ultra-resolution dataset identified large loops previously missed when using Hi-C datasets sequenced at lower depths; therefore, we asked whether large loops have different features from short loops. We first hypothesized that the ability to form large loops is affected by the number of internal extrusion blockage sites, i.e., nested loop anchors. Under this hypothesis, small loops should contain more extrusion-blockage sites compared to large loops. We detect more nested loops inside large loops compared to small loops (Fig. S7M), therefore, the absolute number of potential blockage sites cannot explain why large loops form in some genomic regions. Examining chromatin states, we found that larger loops have a lower proportion of active chromatin and a greater proportion of quiescent chromatin inside the loop (Fig. S7N). These data suggest that chromatin state might impact the progress of loop extrusion; however, we cannot determine if this effect is due to altered extrusion dynamics or different frequencies and positions of cohesin loading and unloading. The correlation between loop size and chromatin status suggests that analysis of loops in low coverage Hi-C datasets biases the analysis toward small loops and therefore active chromatin.

### Supplemental Methods

*Resolution Comparison and Hi-C Quality Metrics*

We used several metrics to evaluate the quality of the full 20.3 billion Hi-C map in comparison to subsampled Hi-C maps (Fig. S6-S7). First, as a simple Boolean metric, the number of bin-pairs with at least one read was plotted as a fraction of the total number of possible bin-pairs. This was done for bin-pairs within a 1 Mb distance and for all intra-chromosomal bin-pairs.  Non-mappable regions were excluded from analysis and identified by searching for rows and columns within the Hi-C matrix that had no mappable read-pairs. Second, the noise estimates were calculated by taking the average of the autocorrelation function (acf), using a lag of 1, for each row within the matrix of distance normalized Hi-C read-pairs. Noise values were then estimated by (acf -1)*-1. For the comparison between the 33.3 billion Hi-C map and subsampled maps, we calculated the acf on a representative region of the matrix extending between chromosome 1: 1 Mb – 10 mb (shown in Fig. S6). We provide a script for noise estimation at https://github.com/JRowleyLab/HiCNoiseMeasurer.

UMAP clustering was performed using DNase, H2AZ, H3K27ac, H3K27me3, H3K36me3, H3K4me1, H3K4me3, H3K9ac, and H3K9me3 obtained by Avocado[5]. AA/AB clustering scores were obtained by taking each point in the A compartment and summing the UMAP cluster distances to the nearest 10 other points labeled as A and dividing by the sum of distances to the nearest 10 other points labeled B. As an alternative we calculated the 5, 10, 50, 100, 500, and 1000 nearest neighbors using the ball tree algorithm in scikit-learn[6] and calculated the average number of neighbors that had opposite compartmental status.

*CTCF Loop Domains*

Domains were identified by the Arrowhead program[7].

### Supplemental Figures

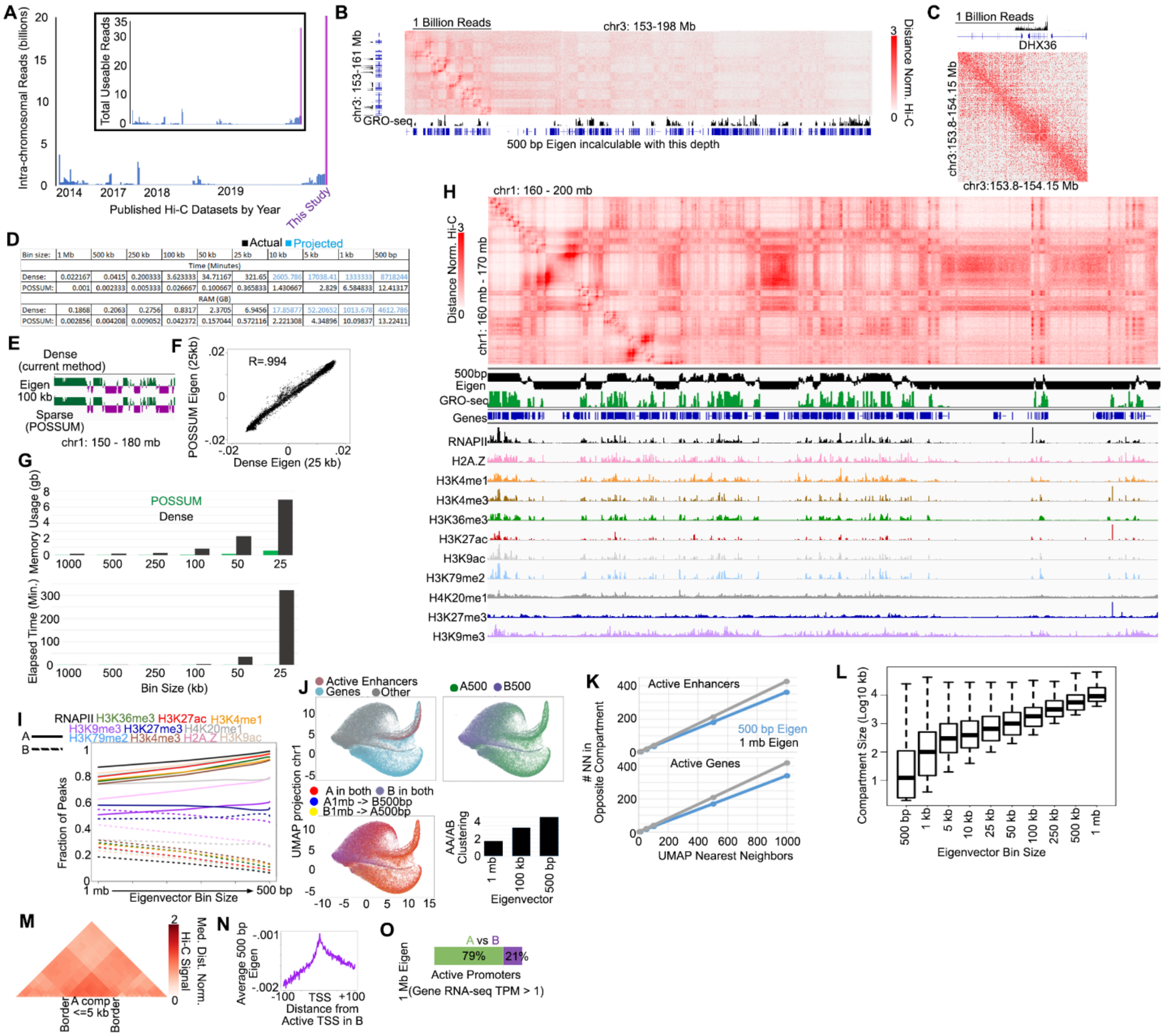


### Supplemental Figure S1

A) Number of intra-chromosomal and total (inset) useable read-pairs in the full LCL Hi-C map (purple) compared to publicly available maps (blue).

B&C) Example of long-range compartment interactions (B) and a compartment domain (C) in a Hi-C map when with only 1 billion intra-chromosomal read-pairs. Black track displays transcription measured by GRO-seq.

D) Table of time and memory usage taken by the current method (dense) vs. POSSUM (sparse) of calculating compartments. Note, that resolutions beyond 25 kb were not possible to calculate using the current method, thus projected values are calculated from a trend line fit to the available datapoints.

E) Eigenvector tracks showing the similarity between the eigenvector called on dense vs sparse (POSSUM) matrices. Comparison of memory usage and time is also shown.

F) Comparison of the eigenvector calculated by current method (dense) vs. POSSUM (sparse) in 25 kb bins. R=.994 (Pearson).

G) Comparison of observed memory usage and time taken by POSSUM (green) vs the current method.

H) Example of compartments seen by distance normalized Hi-C, the eigenvector at 500 bp resolution, and the corresponding transcription (GRO-seq), RNA polymerase (RNAPII), and various histone marks.

I) Fraction of RNA polymerase peaks, or various histone modifications ChIP-seq peaks that were assigned to the A (solid) or B (dashed) compartment when identified with various bin sizes.

J) UMAP clustering of bins based on histone marks. Top Left: Active enhancers (H3K27ac) and active genes (TPM >= 1) are colored orange and blue respectively, other loci are in grey. Bottom Left: Same clustering, but color coded according to compartmental status in the 500 bp or 1 Mb eigenvector. Large amount of orange is due to yellow (reassigned to A) overlapping red (consistently A). Top Right: Same clustering, cut color coded for the A (green) or B (purple) compartmental status at 500 bp resolution. Bottom Right: Clustering distance of each A point to the nearest 10 A points vs the nearest 10 B points for the 500 bp, 100 kb, and 1 Mb compartment eigenvectors.

K) In UMAP plots clustered by histone modification, the number of nearest neighbors which were in opposite compartment calls (Y-axis) when examining various numbers of nearest neighbors (X-axis).

L) Compartment sizes when identified at various resolutions.

M) Median distance normalized Hi-C signal near the diagonal for small, <= 5 kb A compartment domains identified by the POSSUM eigenvector.

N) Average POSSUM eigenvector around TSSs assigned to the B compartment.

O) Percentage of active promoters assigned to A (green) or B (purple) when compartments are identified at 1 Mb resolution.

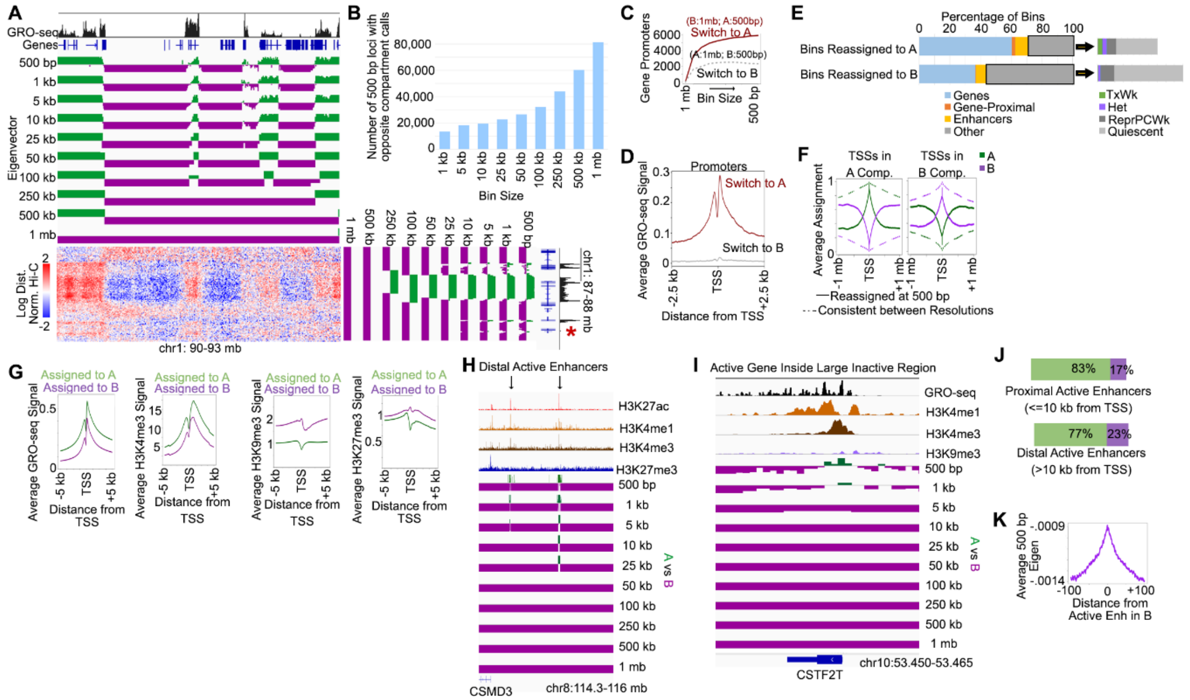


**Supplemental Figure S2**

A) Example of small compartment domains only identifiable at high-resolution (red asterisks). Log transformed and distance normalized Hi-C map is shown alongside the eigenvector tracks at various bin sizes.

B) Number of loci with opposite compartment calls when they are identified in coarser bins.

C) Number of promoters that are reassigned to A (red) or B (grey) compartments when the eigenvector is used at various resolutions.

D) Average GRO-seq signal near TSSs that are reassigned to A (red) of B (grey) compartments.

E) Bins reassigned to A (top) or B (bottom) when using the 500 bp eigenvector and the percentage that overlap with various features.

F) The average A (green) and B (purple) compartmental status at 500 bp resolution for TSSs and the surrounding region for those TSSs reassigned compartments (solid line) vs those consistent between resolutions (dashed).

G) Average GRO-seq, and H3K4me3, H3K9me3, and H3K27me3 ChIP-seq (from left to right) at active gene promoters assigned to A (green) or B (purple) compartments at 500 bp resolution.

H) Examples of distal active enhancers with mismatched compartment assignments at coarse resolutions.

I) Example of an active promoter with mismatched compartment assignment at coarse resolutions.

J) Percentage of active proximal (top) or distal (bottom) enhancers assigned to A (green) or B (purple) when compartments are identified at 1 Mb resolution.

K) Average POSSUM eigenvector around active enhancers assigned to the B compartment.

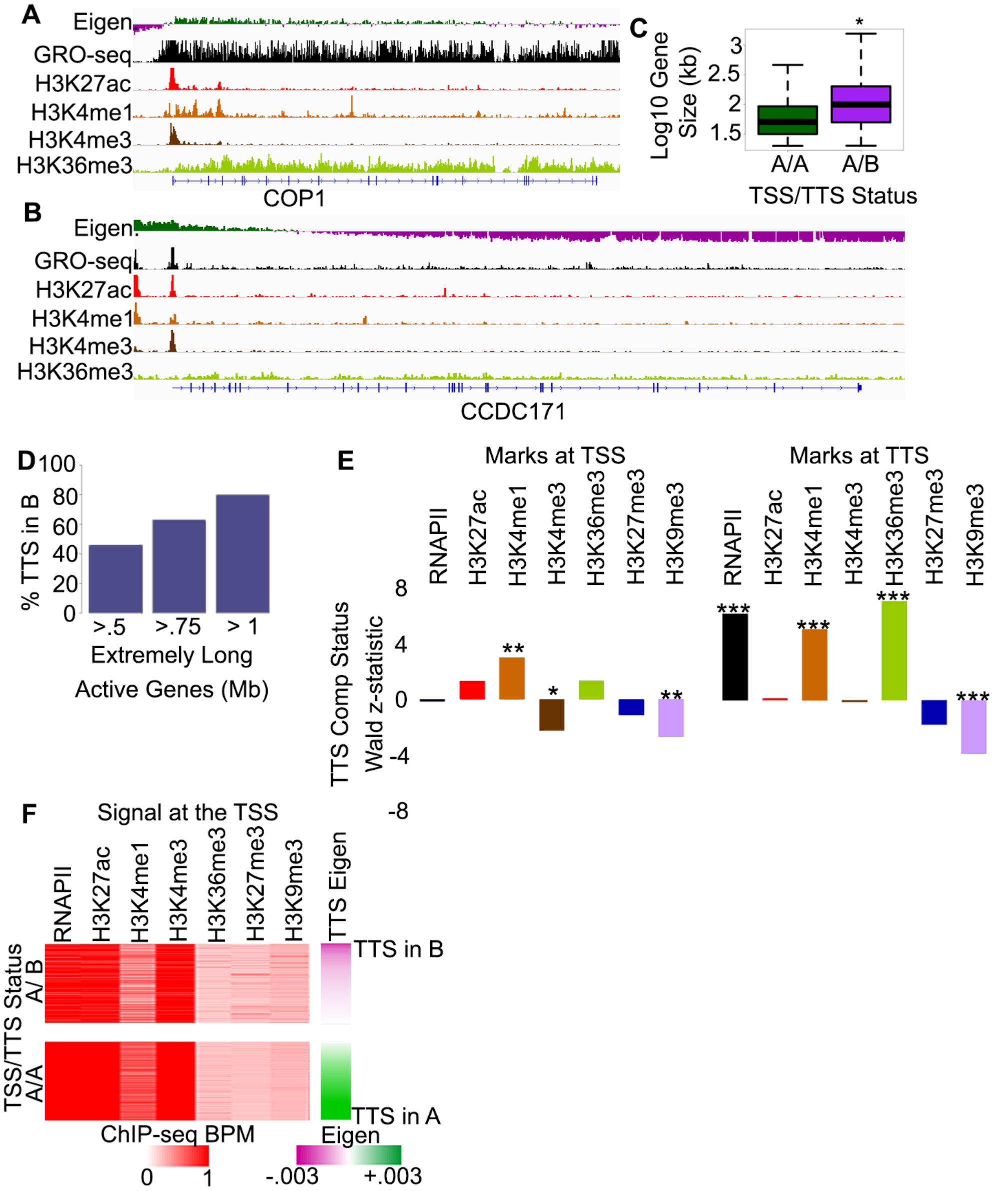


**Supplemental Figure 3**

A&B) Examples of gene with different GRO-seq distributions and different A compartmental distributions along the gene body.

C) Gene size vs. the compartmental status of the TTS. * denotes p-value < .05 Wilcoxon Rank Sum test.

D) Percentage of TTSs assigned to the B compartment for extremely large genes.

E) Results of logistic regression tests for the relationship between marks at the TSS or TTS vs the compartmental status of the TTS. * p<.05, ** p<.01, *** p<.001.

F) ChIP-seq signal at the TSS of discordant A/B genes vs. concordant A/A genes. Genes are sorted by the TTS compartmental signal.

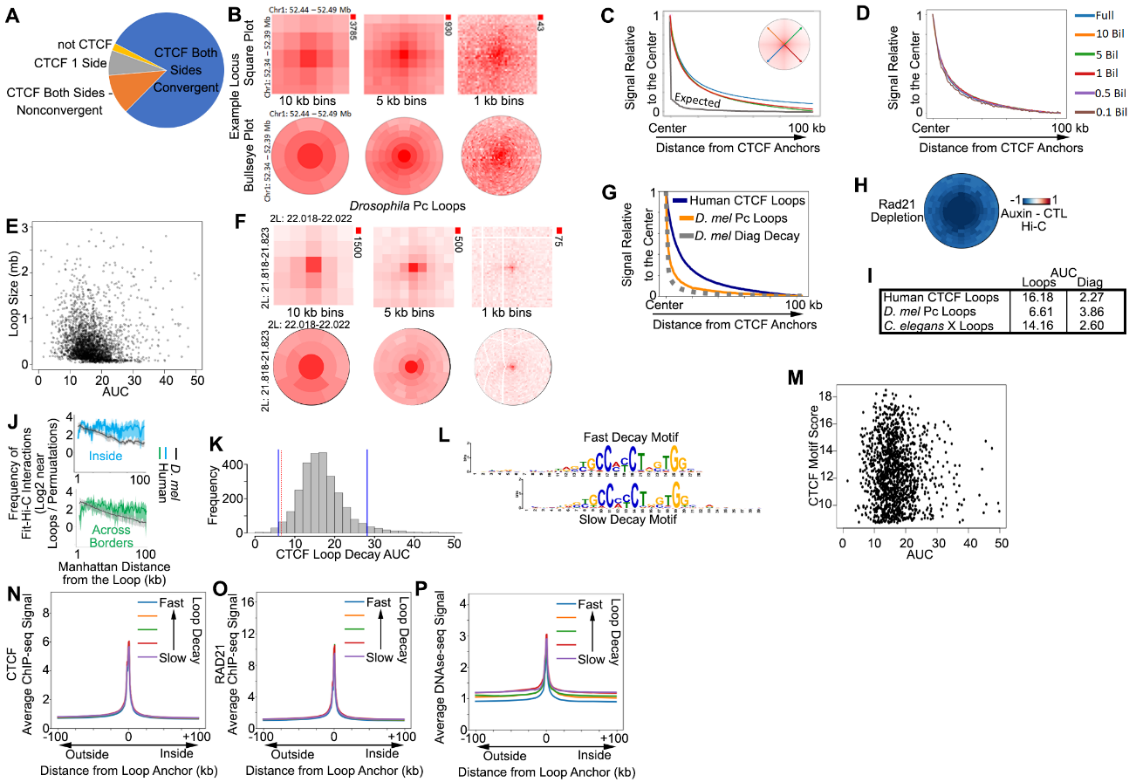


**Supplemental Figure S4**

A) The distribution of loops which had CTCF in convergent orientation on both anchors (blue), in other orientations (orange), on only one side (grey), or had no evidence of CTCF (yellow).

B) Square (top) and bullseye (bottom) views of an example CTCF loop in human cells when binned at 10, 5, or 1 kb.

C) Average decay of signal starting at loops and moving inside (blue), parallel (red and orange), or outward (green) Average signal decaying at the diagonal is shown for reference (grey).

D) Average loop-decay in subsampled maps.

E) The area under the curve (AUC) for the loop signal decay vs size of each CTCF loop.

F) Square (top) and bullseye (bottom) views of an example Pc loop in *D. melanogaster* cells when binned at 10, 5, or 1 kb.

G) Average Hi-C signal in 1 kb bins at each radial distance away from human CTCF loop anchors (blue) vs. *D. melanogaster* Pc loops (orange). Average signal at the *D. melanogaster* Hi-C diagonal is shown for reference (grey).

H) Average plots after Rad21 degradation showing the change in Hi-C signal at the loop as well as in proximal regions.

I) Table of AUC values for loops and the diagonal in human, *D. melanogaster*, and *C. elegans* Hi-C maps.

J) Frequency of enhancer-promoter interactions determined by Fit-Hi-C in the area proximal to CTCF loops in human cells vs *Drosophila* Pc loops. Interactions that are completely interior to loops vs those that cross over one or both anchors are shown.

K) Histogram of loop-decay AUC (area under the curve) values for CTCF loops. Black vertical lines indicate a standard deviation of 1.5. Red vertical line indicates the average AUC of *D. melanogaster* Pc loops.

L) CTCF motifs found at slow and fast decaying loops.

M) Comparison of CTCF motif score at anchors to the decay AUC.

N-P) Average profile of CTCF ChIP-seq signal, RAD21 ChIP-seq signal, and DNAse-seq signal at the anchors of loops that show different diffusiveness.

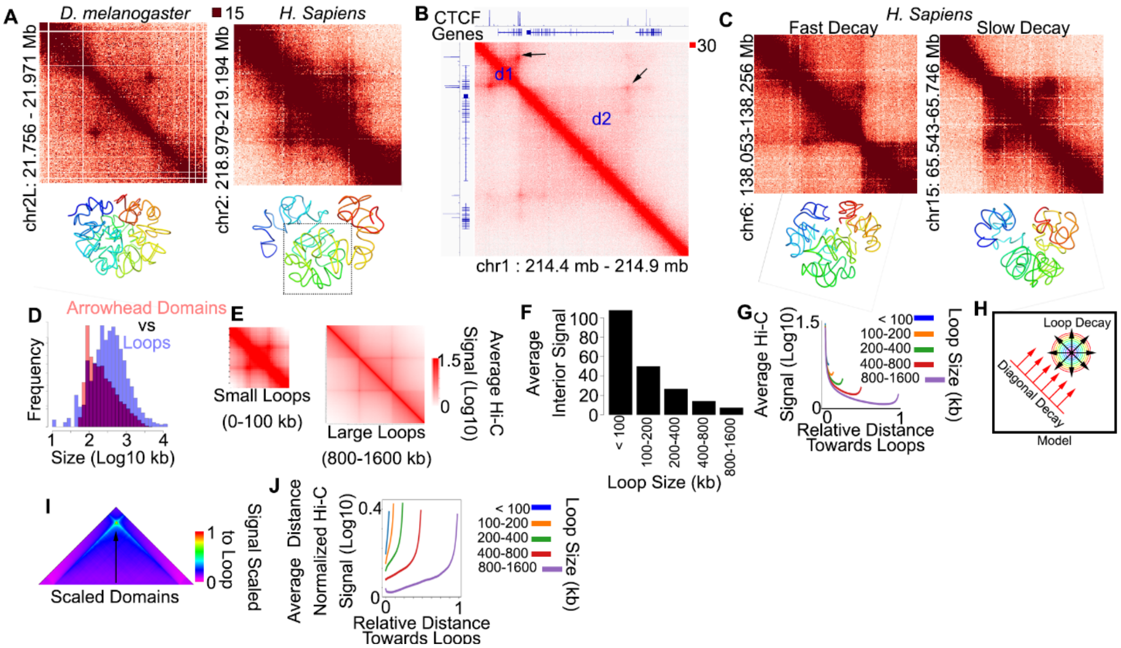


A) Example of a Pc loop in *D. melanogaster* (left) and a CTCF loop in humans (right). Under each Hi-C heat map is the corresponding Pastis model colored by bin progressing blue to red (top left to bottom right).

B) Example of two loops of different sizes and with different interior domain strength. Top track shows CTCF ChIP-seq signal.

C) Example of two same sized CTCF loops with different decay rates. Under each Hi-C heat map is the corresponding Pastis model colored by bin progressing blue to red (top left to bottom right).

D) Histogram of loop sizes (blue) vs the sizes of arrowhead domains (peach).

E) Average Hi-C signal for small vs large loops scaled to the average size in that category.

F) Average Hi-C signal inside of loops of various sizes.

G) Average Hi-C signal at each distance from the diagonal internal to loops. X-axis distance is set relative to the largest loop category.

H) Diagram showing how polymeric distance (diagonal decay) could combine with diffusive CTCF loop signal (loop decay).

I) Average distance normalized signal relative to the punctate loop signal.

J) Average distance normalized Hi-C signal at each distance from the diagonal internal to loops. X-axis distance is set relative to the largest loop category.

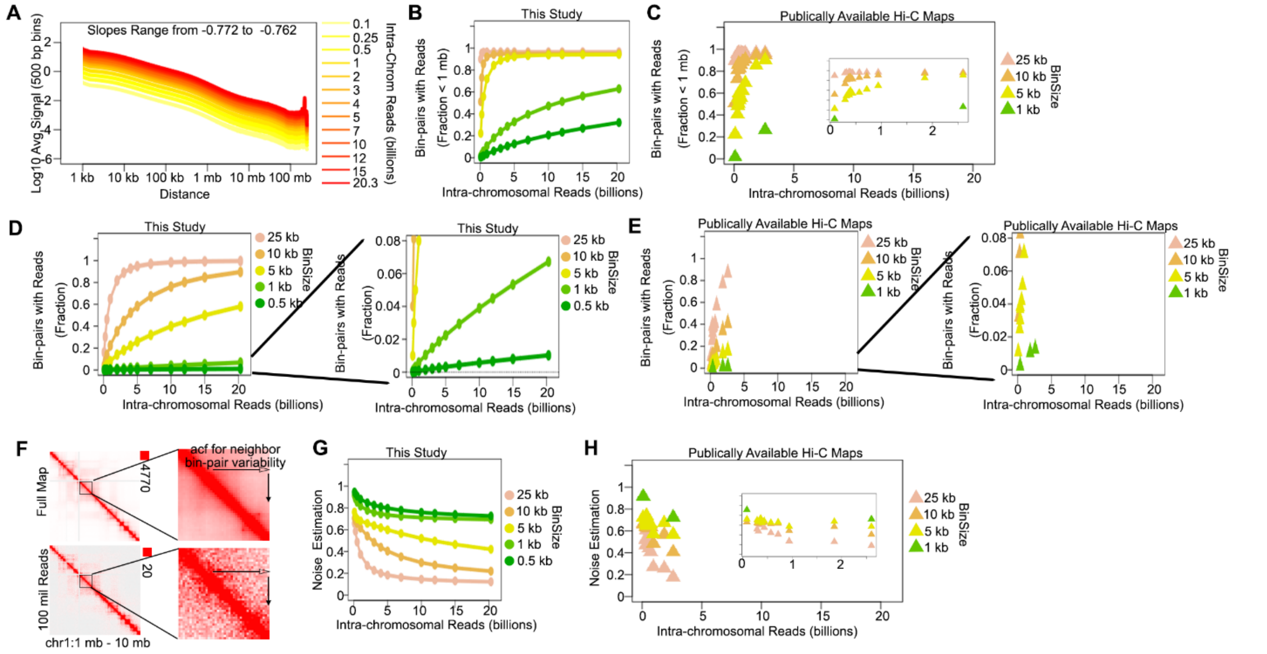


**Supplemental Figure S6**

A) Diagonal decay in our full 20.3 billion map and in maps with subsampled read-pairs.

B&C) Fraction of bin-pairs at distances < 1 Mb that have at least 1 read in the full and subsampled maps (B) and in published maps (C).

D&E) Fraction of bin-pairs at all distances that have at least 1 read considering all distances in the full 20.3 billion map and in maps with subsampled read-pairs (D) or in published maps (E). Right panels show the data when zoomed in on the y-axis.

F) Example of how sequencing coverage reduces noise in Hi-C maps and how the autocorrelation function (acf), which measures similarities between neighboring bin-pairs can be used to estimate noise.

G&H) Noise estimated from the inverse of the autocorrelation function for the full and subsampled maps (G) and in published Hi-C maps (H).

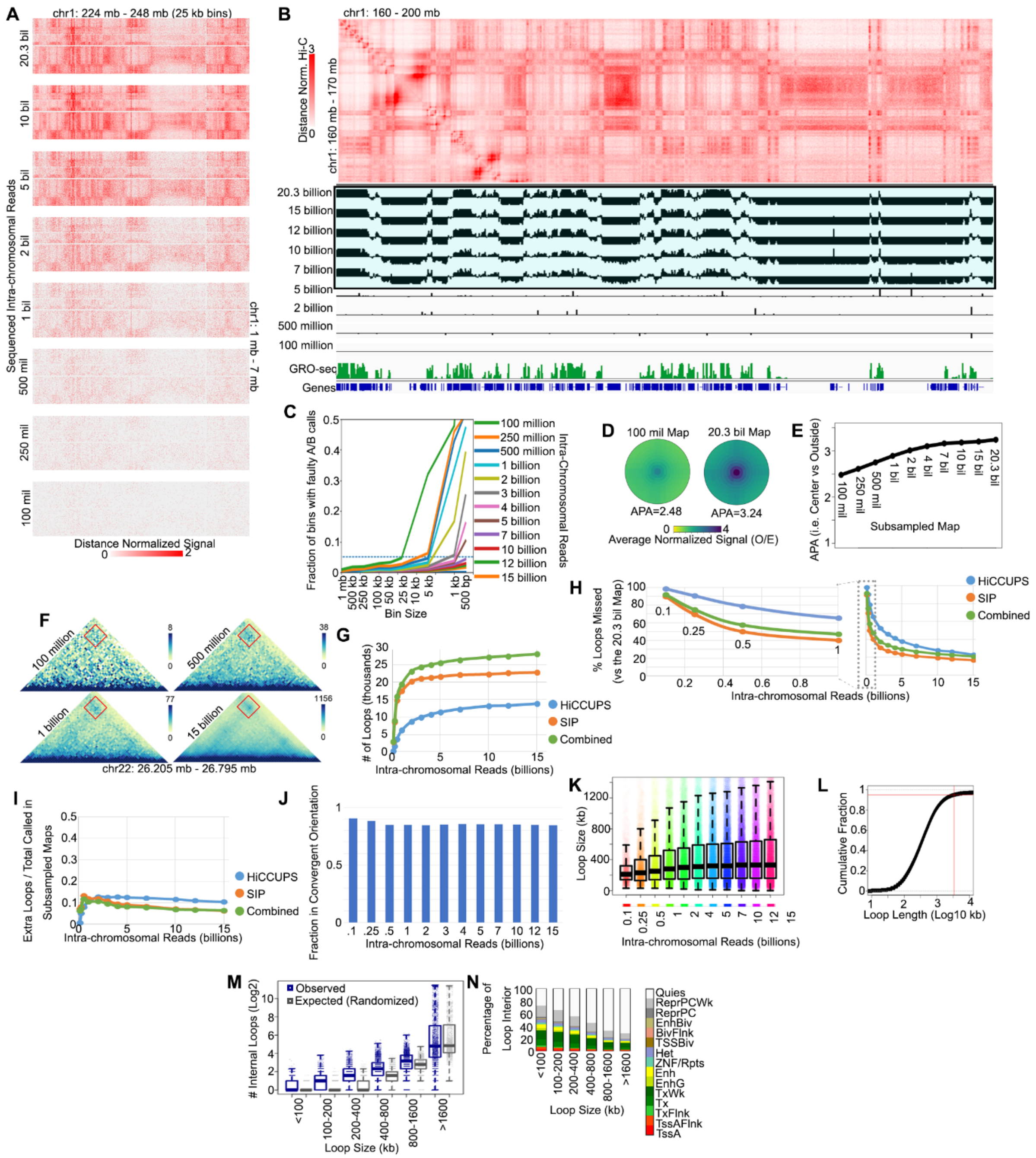


**Supplemental Figure S7**

A) Example of compartments seen by distance normalized Hi-C and the eigenvector at 500 bp resolution in maps with various depth. Active sites are represented by high GRO-seq signal.

B) Comparison of 500 bp compartment identification in maps with various sequencing depths.

C) Fraction of mismatched compartment calls at various resolutions and between maps with various sequencing depths.

D) Average CTCF loops signal in a Hi-C contact map with 100 million reads vs 20.3 billion reads.

E) Aggregate peaks analysis (APA) in subsampled maps and in the full 20.3 billion map. Numbers represent intra-chromosomal read pairs.

F) Example of a loop that is missed with lower sequencing depth. Numbers represent intra-chromosomal read pairs.

G) Number of loops called in subsampled maps as called by HiCCUPS (blue), SIP (orange), or when combined (green).

H) Percentage of loops missed in subsampled maps.

I) The number of loops called only in the subsampled map vs the full 20.3 billion map as a ratio to the total called for HiCCUPS (blue), SIP (orange), or when combined (green).

J) The fraction of loops called in subsampled maps that have CTCF in convergent orientation.

K) Sizes of loops identifiable in Hi-C maps with various sequencing depths.

L) Cumulative fraction of loop sizes in the full map.

M) Number of loops that are internal to those of various sizes (blue) compared to what would be expected by random genomic regions of similar size (grey). * $p<.05$ Wilcoxon Rank Sum test.

N) Percentage of various sized loop interiors that correspond to various chromatin states.

### Supplemental Tables

Sequenced Read Pairs: 42,467,875,383
 Normal Paired: 23,937,977,379 (56.37%)
 Chimeric Paired: 14,294,168,018 (33.66%)
 Chimeric Ambiguous: 3,382,008,102 (7.96%)
 Unmapped: 853,721,882 (2.01%)
 Alignable (Normal+Chimeric Paired): 38,232,145,397 (90.03%)
Unique Reads: 34,806,960,717
PCR Duplicates: 3,385,126,713
Optical Duplicates: 203,288,499
Intra-fragment Reads: 840,557,070 (1.98% / 2.41%)
Below MAPQ Threshold: 3,217,457,396 (7.58% / 9.23%)
Hi-C Contacts: 30,815,303,487 (72.56% / 88.36%)
 Ligation Motif Present: 0  (0.00% / 0.00%)
 3' Bias (Long Range): 0% - 0%
 Pair Type %(L-I-O-R): 25% - 25% - 25% - 25%
Inter-chromosomal: 7,663,533,305  (18.05% / 21.98%)
Intra-chromosomal: 23,151,770,182  (54.52% / 66.39%)
Short Range (<20Kb): 7,905,477,803  (18.62% / 22.67%)
Long Range (>20Kb): 15,246,211,030  (35.90% / 43.72%)

**Supplemental Table S1**. Mapping and Filtering Statistics for the Ultra-Res Hi-C map

| Accession | File Type | | Total reads | Cis reads | Short (<20kb) | Trans | Date Published |
|---|---|---|---|---|---|---|---|
| 4DNES7L8Z2KV | contact | list-combined | 6.94E+08 | 4.10E+08 | 1.28E+08 | 1.55E+08 | 2/21/2017 |
| 4DNES49TDMJM | contact | list-combined | 7.18E+08 | 4.25E+08 | 1.76E+08 | 1.16E+08 | 2/21/2017 |
| 4DNESWEF2AHT | contact | list-combined | 5.79E+08 | 3.87E+08 | 1.01E+08 | 9.11E+07 | 2/21/2017 |
| 4DNESHGZUBL9 | contact | list-combined | 5.64E+08 | 3.35E+08 | 1.20E+08 | 1.09E+08 | 2/21/2017 |
| 4DNES5R3O24W | contact | list-combined | 5.76E+08 | 3.23E+08 | 1.68E+08 | 8.53E+07 | 2/21/2017 |
| 4DNESU4BQU4G | contact | list-combined | 8.12E+08 | 5.32E+08 | 1.55E+08 | 1.26E+08 | 2/21/2017 |
| 4DNES1QUXG92 | contact | list-combined | 4.53E+08 | 1.26E+08 | 2.64E+08 | 6.29E+07 | 3/20/2017 |
| 4DNESWDLDMGN | contact | list-combined | 5.14E+08 | 2.68E+08 | 1.70E+08 | 7.61E+07 | 3/20/2017 |
| 4DNESQMM4EBN | contact | list-combined | 4.11E+08 | 1.38E+08 | 2.33E+08 | 4.08E+07 | 3/20/2017 |
| 4DNESKKSKG7Y | contact | list-combined | 1.07E+09 | 6.06E+08 | 3.29E+08 | 1.38E+08 | 3/20/2017 |
| 4DNES76KXUJ3 | contact | list-combined | 7.08E+08 | 4.84E+08 | 1.36E+08 | 8.81E+07 | 5/4/2017 |
| 4DNESNLXBFMY | contact | list-combined | 4.16E+08 | 2.46E+08 | 9.28E+07 | 7.73E+07 | 5/4/2017 |
| 4DNESIU6F8HF | contact | list-combined | 4.23E+08 | 2.71E+08 | 9.12E+07 | 6.13E+07 | 5/4/2017 |
| 4DNESSQU7B76 | contact | list-combined | 4.14E+08 | 2.57E+08 | 9.46E+07 | 6.21E+07 | 5/4/2017 |
| 4DNES7RYT7KA | contact | list-combined | 6.26E+07 | 3.53E+07 | 1.66E+07 | 1.06E+07 | 7/12/2017 |
| 4DNES6U3U9PJ | contact | list-combined | 1.49E+08 | 8.40E+07 | 3.71E+07 | 2.81E+07 | 7/12/2017 |
| 4DNESTHQ7CR1 | contact | list-combined | 6.53E+07 | 3.53E+07 | 1.80E+07 | 1.19E+07 | 7/12/2017 |
| 4DNESO6J5SH9 | contact | list-combined | 1.16E+08 | 5.58E+07 | 3.30E+07 | 2.69E+07 | 7/12/2017 |
| 4DNESUQT299T | contact | list-combined | 6.20E+07 | 3.44E+07 | 1.68E+07 | 1.08E+07 | 7/12/2017 |
| 4DNESMXBLGKA | contact | list-combined | 1.45E+08 | 9.20E+07 | 3.12E+07 | 2.20E+07 | 7/12/2017 |
| 4DNESSL62JQK | contact | list-combined | 2.03E+08 | 1.14E+08 | 4.78E+07 | 4.09E+07 | 7/12/2017 |
| 4DNESWPY49OL | contact | list-combined | 2.48E+08 | 1.41E+08 | 7.42E+07 | 3.28E+07 | 7/12/2017 |
| 4DNESXZHIUAU | contact | list-combined | 2.75E+08 | 1.29E+08 | 7.74E+07 | 6.84E+07 | 7/12/2017 |
| 4DNESICXDTH7 | contact | list-combined | 1.89E+08 | 9.12E+07 | 5.69E+07 | 4.04E+07 | 7/12/2017 |
| 4DNES1J8MC4Q | contact | list-combined | 1.99E+08 | 9.40E+07 | 6.16E+07 | 4.30E+07 | 7/12/2017 |
| 4DNES3JWOSVS | contact | list-combined | 1.05E+08 | 5.12E+07 | 3.30E+07 | 2.08E+07 | 7/12/2017 |
| 4DNESOBRQ1WW | contact | list-combined | 6.66E+07 | 4.76E+07 | 1.70E+07 | 2.06E+06 | 7/12/2017 |
| 4DNESTRKO6LB | contact | list-combined | 8.69E+07 | 2.16E+07 | 3.30E+07 | 3.23E+07 | 7/12/2017 |
| 4DNES4KDIQNN | contact | list-combined | 5.73E+08 | 2.59E+08 | 1.55E+08 | 1.59E+08 | 10/5/2017 |
| 4DNES7RDRS69 | contact | list-combined | 6.16E+08 | 2.84E+08 | 1.58E+08 | 1.75E+08 | 10/5/2017 |
| 4DNESCOJ3ADI | contact | list-combined | 6.27E+08 | 3.04E+08 | 1.53E+08 | 1.71E+08 | 10/5/2017 |
| 4DNESJV9TH8Q | contact | list-combined | 2.55E+09 | 1.24E+09 | 4.39E+08 | 8.67E+08 | 10/5/2017 |
| 4DNES3QAGOZZ | contact | list-combined | 2.69E+09 | 1.38E+09 | 5.36E+08 | 7.73E+08 | 10/5/2017 |
| 4DNESD117NVX | contact | list-combined | 3.20E+07 | 1.64E+07 | 6.76E+06 | 8.86E+06 | 10/5/2017 |
| 4DNESLLMJ3JO | contact | list-combined | 2.76E+07 | 1.35E+07 | 6.50E+06 | 7.65E+06 | 10/5/2017 |
| 4DNES64ITOWM | contact | list-combined | 2.49E+07 | 1.30E+07 | 5.58E+06 | 6.26E+06 | 10/5/2017 |
| 4DNESNGDNQSG | contact | list-combined | 1.88E+07 | 8.86E+06 | 3.92E+06 | 6.06E+06 | 10/5/2017 |
| 4DNESD6N2Y6L | contact | list-combined | 9.11E+06 | 4.08E+06 | 2.24E+06 | 2.79E+06 | 10/5/2017 |
| 4DNESFPM1OFO | contact | list-combined | 8.59E+06 | 4.24E+06 | 1.26E+06 | 3.09E+06 | 10/5/2017 |
| 4DNES7N8J8KG | contact | list-combined | 1.06E+07 | 5.30E+06 | 1.87E+06 | 3.43E+06 | 10/5/2017 |

| | | | | | | | |
|---|---|---|---|---|---|---|---|
| 4DNESOGIV7NG | contact | list-combined | 6.45E+08 | 3.31E+08 | 1.77E+08 | 1.38E+08 | 10/5/2017 |
| 4DNESOQTEWX8 | contact | list-combined | 1.61E+08 | 8.01E+07 | 3.76E+07 | 4.30E+07 | 10/5/2017 |
| 4DNESMLKX1ZM | contact | list-combined | 1.74E+08 | 7.60E+07 | 4.60E+07 | 5.23E+07 | 10/5/2017 |
| 4DNES1NX2TKW | contact | list-combined | 3.68E+08 | 1.51E+08 | 1.75E+08 | 4.15E+07 | 10/19/2017 |
| 4DNESQT4SR5G | contact | list-combined | 3.90E+08 | 1.55E+08 | 1.92E+08 | 4.35E+07 | 10/19/2017 |
| 4DNES3Y26CEU | contact | list-combined | 3.77E+08 | 1.94E+08 | 1.34E+08 | 4.90E+07 | 10/19/2017 |
| 4DNESUUFHGKJ | contact | list-combined | 3.95E+08 | 1.53E+08 | 1.99E+08 | 4.34E+07 | 10/19/2017 |
| 4DNESGC3Z7E3 | contact | list-combined | 3.61E+08 | 1.50E+08 | 1.73E+08 | 3.83E+07 | 10/19/2017 |
| 4DNES68MSHVU | contact | list-combined | 3.99E+08 | 1.57E+08 | 1.97E+08 | 4.45E+07 | 10/19/2017 |
| 4DNESXS1M9JR | contact | list-combined | 3.68E+08 | 1.67E+08 | 1.56E+08 | 4.50E+07 | 10/19/2017 |
| 4DNESBBYGJFA | contact | list-combined | 3.96E+08 | 1.60E+08 | 1.91E+08 | 4.56E+07 | 10/19/2017 |
| 4DNESU4Y9CBF | contact | list-combined | 5.52E+07 | 3.42E+07 | 1.21E+07 | 8.86E+06 | 10/19/2017 |
| 4DNESXI5NKKT | contact | list-combined | 6.39E+07 | 4.00E+07 | 1.33E+07 | 1.05E+07 | 10/19/2017 |
| 4DNESUCLJAZ8 | contact | list-combined | 1.24E+08 | 8.15E+07 | 2.68E+07 | 1.57E+07 | 10/19/2017 |
| 4DNESJ9SIAV5 | contact | list-combined | 5.37E+09 | 2.82E+09 | 1.05E+09 | 1.51E+09 | 10/19/2017 |
| 4DNESDXUWBD9 | contact | list-combined | 4.26E+09 | 2.11E+09 | 1.52E+09 | 6.27E+08 | 10/19/2017 |
| 4DNESI9RVI9Y | contact | list-combined | 1.04E+07 | 4.86E+06 | 1.89E+06 | 3.60E+06 | 4/26/2018 |
| 4DNESJ5Y5ZN8 | contact | list-combined | 1.25E+07 | 7.41E+06 | 2.75E+06 | 2.34E+06 | 4/26/2018 |
| 4DNESKZ1WYCN | contact | list-combined | 1.14E+07 | 6.46E+06 | 2.75E+06 | 2.17E+06 | 4/26/2018 |
| 4DNESKY5IGMT | contact | list-combined | 7.37E+06 | 4.23E+06 | 1.51E+06 | 1.63E+06 | 4/26/2018 |
| 4DNESWSTOQ92 | contact | list-combined | 8.20E+06 | 3.35E+06 | 1.66E+06 | 3.19E+06 | 4/26/2018 |
| 4DNES51DMKBB | contact | list-combined | 9.11E+06 | 4.66E+06 | 2.02E+06 | 2.42E+06 | 4/26/2018 |
| 4DNESHDMG1LU | contact | list-combined | 1.02E+07 | 4.94E+06 | 2.76E+06 | 2.55E+06 | 4/26/2018 |
| 4DNESTHU1G8W | contact | list-combined | 1.11E+07 | 5.45E+06 | 2.13E+06 | 3.56E+06 | 4/26/2018 |
| 4DNESG1W727L | contact | list-combined | 6.34E+06 | 3.12E+06 | 1.15E+06 | 2.07E+06 | 4/26/2018 |
| 4DNESHN4FT5O | contact | list-combined | 6.91E+06 | 3.27E+06 | 2.01E+06 | 1.63E+06 | 4/26/2018 |
| 4DNESD21O5UU | contact | list-combined | 1.02E+07 | 5.26E+06 | 2.11E+06 | 2.88E+06 | 4/26/2018 |
| 4DNES1XC6D9N | contact | list-combined | 1.39E+07 | 7.38E+06 | 3.19E+06 | 3.34E+06 | 4/26/2018 |
| 4DNESZ1UN35D | contact | list-combined | 1.02E+07 | 5.08E+06 | 2.79E+06 | 2.33E+06 | 4/26/2018 |
| 4DNESS3E2ITC | contact | list-combined | 9.30E+06 | 5.00E+06 | 1.81E+06 | 2.49E+06 | 4/26/2018 |
| 4DNESCQZYHLJ | contact | list-combined | 3.52E+07 | 1.92E+07 | 7.89E+06 | 8.08E+06 | 4/26/2018 |
| 4DNESEZ64E3W | contact | list-combined | 6.41E+06 | 2.98E+06 | 1.83E+06 | 1.60E+06 | 4/26/2018 |
| 4DNESM6U2PMJ | contact | list-combined | 8.00E+06 | 4.23E+06 | 1.40E+06 | 2.37E+06 | 4/26/2018 |
| 4DNESFSFD7PV | contact | list-combined | 1.11E+07 | 5.73E+06 | 2.61E+06 | 2.78E+06 | 4/26/2018 |
| 4DNESH3WR89R | contact | list-combined | 9.20E+06 | 4.82E+06 | 2.06E+06 | 2.33E+06 | 4/26/2018 |
| 4DNESIXD38O6 | contact | list-combined | 6.69E+06 | 3.46E+06 | 1.48E+06 | 1.75E+06 | 4/26/2018 |
| 4DNESUZCBCG9 | contact | list-combined | 3.21E+07 | 1.73E+07 | 6.96E+06 | 7.85E+06 | 4/26/2018 |
| 4DNESFJ1LEZ2 | contact | list-combined | 1.54E+07 | 8.45E+06 | 3.83E+06 | 3.09E+06 | 4/26/2018 |
| 4DNES5S3N3UN | contact | list-combined | 9.45E+06 | 4.86E+06 | 1.55E+06 | 3.04E+06 | 4/26/2018 |
| 4DNES2Z1IO3Y | contact | list-combined | 1.24E+07 | 7.04E+06 | 2.29E+06 | 3.04E+06 | 4/26/2018 |
| 4DNESA51WBOJ | contact | list-combined | 1.74E+07 | 8.07E+06 | 2.27E+06 | 7.07E+06 | 4/26/2018 |

| | | | | | | | |
|---|---|---|---|---|---|---|---|
| 4DNESV19UH2A | contact | list-combined | 2.05E+07 | 1.02E+07 | 3.83E+06 | 6.45E+06 | 4/26/2018 |
| 4DNESXWCEU6I | contact | list-combined | 6.85E+06 | 3.24E+06 | 1.83E+06 | 1.78E+06 | 4/26/2018 |
| 4DNESP7EFMLT | contact | list-combined | 1.21E+07 | 6.29E+06 | 2.64E+06 | 3.15E+06 | 4/26/2018 |
| 4DNESQNC15UK | contact | list-combined | 9.49E+06 | 5.03E+06 | 1.83E+06 | 2.63E+06 | 4/26/2018 |
| 4DNESZRLWUML | contact | list-combined | 1.09E+07 | 5.52E+06 | 2.29E+06 | 3.06E+06 | 4/26/2018 |
| 4DNES84ZICUS | contact | list-combined | 7.01E+06 | 3.57E+06 | 1.59E+06 | 1.85E+06 | 4/26/2018 |
| 4DNESK7WOJRD | contact | list-combined | 2.98E+07 | 1.56E+07 | 6.29E+06 | 7.96E+06 | 4/26/2018 |
| 4DNESLFAFVGO | contact | list-combined | 7.18E+06 | 3.48E+06 | 1.85E+06 | 1.86E+06 | 4/26/2018 |
| 4DNESOSF88NZ | contact | list-combined | 5.76E+06 | 2.92E+06 | 1.55E+06 | 1.28E+06 | 4/26/2018 |
| 4DNESXVOAC79 | contact | list-combined | 1.10E+07 | 5.61E+06 | 2.25E+06 | 3.09E+06 | 4/26/2018 |
| 4DNESD7DATO7 | contact | list-combined | 9.10E+06 | 4.78E+06 | 1.66E+06 | 2.66E+06 | 4/26/2018 |
| 4DNES2N7SYC4 | contact | list-combined | 1.38E+07 | 7.25E+06 | 3.71E+06 | 2.85E+06 | 4/26/2018 |
| 4DNES8YBULP1 | contact | list-combined | 7.98E+06 | 4.28E+06 | 1.60E+06 | 2.10E+06 | 4/26/2018 |
| 4DNESFNBTRO3 | contact | list-combined | 3.16E+07 | 1.75E+07 | 6.78E+06 | 7.37E+06 | 4/26/2018 |
| 4DNESDC4XDHT | contact | list-combined | 9.71E+06 | 4.79E+06 | 2.21E+06 | 2.71E+06 | 4/26/2018 |
| 4DNESERXQSXY | contact | list-combined | 3.19E+06 | 1.52E+06 | 7.25E+05 | 9.46E+05 | 4/26/2018 |
| 4DNESHF65RC4 | contact | list-combined | 3.19E+07 | 1.62E+07 | 6.97E+06 | 8.76E+06 | 4/26/2018 |
| 4DNESKLFZ31S | contact | list-combined | 3.68E+07 | 1.92E+07 | 8.04E+06 | 9.58E+06 | 4/26/2018 |
| 4DNES4GDH4BG | contact | list-combined | 1.16E+09 | 5.26E+08 | 3.58E+08 | 2.79E+08 | 4/26/2018 |
| 4DNES7C6LBWI | contact | list-combined | 9.15E+08 | 5.06E+08 | 2.19E+08 | 1.91E+08 | 4/26/2018 |
| 4DNES3O1B45O | contact | list-combined | 3.40E+08 | 1.73E+08 | 8.43E+07 | 8.29E+07 | 4/26/2018 |
| 4DNESE1VMAMD | contact | list-combined | 3.84E+08 | 2.25E+08 | 1.01E+08 | 5.77E+07 | 4/26/2018 |
| 4DNES25ABNZ1 | contact | list-combined | 1.87E+09 | 1.10E+09 | 2.82E+08 | 4.92E+08 | 4/26/2018 |
| 4DNES7ODZ4MZ | contact | list-combined | 1.20E+09 | 6.86E+08 | 2.13E+08 | 3.05E+08 | 4/26/2018 |
| 4DNESYX7AQRY | contact | list-combined | 1.13E+09 | 6.16E+08 | 2.46E+08 | 2.63E+08 | 4/26/2018 |
| 4DNESVZ6QH33 | contact | list-combined | 2.39E+08 | 1.43E+08 | 5.53E+07 | 4.06E+07 | 7/16/2019 |
| 4DNESR93N4E3 | contact | list-combined | 2.09E+08 | 1.28E+08 | 4.01E+07 | 4.11E+07 | 7/16/2019 |
| 4DNES9J6QJQS | contact | list-combined | 3.05E+08 | 1.64E+08 | 6.01E+07 | 8.11E+07 | 7/16/2019 |
| 4DNESQICH2XW | contact | list-combined | 5.18E+08 | 9.84E+07 | 9.00E+07 | 3.29E+08 | 7/16/2019 |
| 4DNES1E3ET5M | contact | list-combined | 1.20E+08 | 5.44E+07 | 2.47E+07 | 4.08E+07 | 7/16/2019 |
| 4DNESQV6Y4JL | contact | list-combined | 4.60E+08 | 1.86E+08 | 8.60E+07 | 1.88E+08 | 7/16/2019 |
| 4DNESFM66XDL | contact | list-combined | 3.18E+08 | 1.44E+08 | 3.81E+07 | 1.36E+08 | 7/16/2019 |
| 4DNESZUWCRVN | contact | list-combined | 1.24E+08 | 4.37E+07 | 1.39E+07 | 6.60E+07 | 7/16/2019 |
| 4DNESCQRIZ7D | contact | list-combined | 3.67E+08 | 1.13E+08 | 3.66E+07 | 2.18E+08 | 7/16/2019 |
| 4DNESLVTLWPX | contact | list-combined | 3.83E+08 | 8.71E+07 | 3.76E+07 | 2.59E+08 | 7/16/2019 |
| 4DNESF17LNZE | contact | list-combined | 6.02E+08 | 1.21E+08 | 8.31E+07 | 3.98E+08 | 7/16/2019 |
| 4DNESSH9ICEW | contact | list-combined | 4.07E+08 | 4.28E+07 | 3.69E+07 | 3.28E+08 | 7/16/2019 |
| 4DNESC23ZYOF | contact | list-combined | 1.87E+08 | 7.73E+07 | 2.17E+07 | 8.80E+07 | 7/16/2019 |
| 4DNES54GS5KI | contact | list-combined | 2.29E+08 | 9.26E+07 | 2.29E+07 | 1.13E+08 | 7/16/2019 |
| 4DNESPXSO8GB | contact | list-combined | 3.03E+07 | 1.01E+07 | 3.49E+06 | 1.67E+07 | 7/16/2019 |
| 4DNESHJVC7MP | contact | list-combined | 2.43E+07 | 1.01E+07 | 3.21E+06 | 1.10E+07 | 7/16/2019 |

| | | | | | | | |
|---|---|---|---|---|---|---|---|
| 4DNESMG7JML8 | contact | list-combined | 2.66E+07 | 7.19E+06 | 2.95E+06 | 1.64E+07 | 7/16/2019 |
| 4DNESYVC49SD | contact | list-combined | 3.18E+07 | 3.27E+06 | 3.28E+06 | 2.52E+07 | 7/16/2019 |
| 4DNEST7Y7S69 | contact | list-combined | 1.83E+08 | 3.07E+07 | 1.93E+07 | 1.33E+08 | 7/16/2019 |
| 4DNESFOADERB | contact | list-combined | 1.19E+08 | 5.46E+07 | 5.22E+07 | 1.24E+07 | 9/10/2019 |
| 4DNESFI64TG3 | contact | list-combined | 1.31E+08 | 6.41E+07 | 5.00E+07 | 1.65E+07 | 9/10/2019 |
| 4DNESU95RUNO | contact | list-combined | 1.09E+08 | 6.09E+07 | 2.26E+07 | 2.59E+07 | 9/10/2019 |
| 4DNESPOC41XG | contact | list-combined | 1.09E+08 | 6.15E+07 | 2.33E+07 | 2.44E+07 | 9/10/2019 |
| 4DNES9BTO2FB | contact | list-combined | 8.46E+07 | 3.43E+07 | 1.22E+07 | 3.82E+07 | 11/4/2019 |
| 4DNESVR7L225 | contact | list-combined | 7.36E+07 | 3.94E+07 | 9.72E+06 | 2.44E+07 | 11/4/2019 |
| 4DNESCJHJ477 | contact | list-combined | 6.91E+07 | 3.73E+07 | 9.89E+06 | 2.19E+07 | 11/4/2019 |
| 4DNESS4NRI4U | contact | list-combined | 5.41E+07 | 2.91E+07 | 7.17E+06 | 1.77E+07 | 11/4/2019 |
| 4DNESPLDAR9W | contact | list-combined | 5.61E+07 | 2.89E+07 | 8.00E+06 | 1.92E+07 | 11/4/2019 |
| 4DNESI74G82F | contact | list-combined | 6.63E+07 | 3.44E+07 | 9.67E+06 | 2.22E+07 | 11/4/2019 |
| 4DNESXSGHXDQ | contact | list-combined | 6.13E+07 | 3.08E+07 | 8.29E+06 | 2.22E+07 | 11/4/2019 |
| 4DNESAEUDBTF | contact | list-combined | 9.80E+07 | 4.81E+07 | 1.36E+07 | 3.63E+07 | 11/4/2019 |
| 4DNESGQ1XP78 | contact | list-combined | 1.00E+08 | 4.71E+07 | 1.50E+07 | 3.79E+07 | 11/4/2019 |
| 4DNESYJGWM4L | contact | list-combined | 5.47E+07 | 2.81E+07 | 7.70E+06 | 1.89E+07 | 11/4/2019 |
| 4DNES25QLF7A | contact | list-combined | 6.99E+07 | 3.30E+07 | 1.07E+07 | 2.61E+07 | 11/4/2019 |
| 4DNESDRC93UG | contact | list-combined | 7.08E+07 | 3.16E+07 | 1.19E+07 | 2.73E+07 | 11/4/2019 |
| 4DNESAM3IMUG | contact | list-combined | 4.30E+07 | 2.00E+07 | 6.59E+06 | 1.64E+07 | 11/4/2019 |
| 4DNESQ5LDJKC | contact | list-combined | 4.63E+07 | 2.15E+07 | 7.76E+06 | 1.71E+07 | 11/4/2019 |
| 4DNESDW7JTOW | contact | list-combined | 4.96E+07 | 2.29E+07 | 7.65E+06 | 1.91E+07 | 11/4/2019 |
| 4DNES1V3OHQH | contact | list-combined | 5.71E+07 | 2.49E+07 | 9.06E+06 | 2.32E+07 | 11/4/2019 |
| 4DNESALYYHNP | contact | list-combined | 6.74E+07 | 3.06E+07 | 1.06E+07 | 2.62E+07 | 11/4/2019 |
| 4DNESYK6LDN6 | contact | list-combined | 4.13E+07 | 1.67E+07 | 9.20E+06 | 1.54E+07 | 11/4/2019 |
| 4DNESYB59H2X | contact | list-combined | 5.15E+07 | 3.05E+07 | 1.13E+07 | 9.76E+06 | 11/4/2019 |
| 4DNESA2GN9N4 | contact | list-combined | 4.86E+07 | 2.68E+07 | 1.01E+07 | 1.17E+07 | 11/4/2019 |
| 4DNESFMZQ64I | contact | list-combined | 4.74E+07 | 2.55E+07 | 1.03E+07 | 1.17E+07 | 11/4/2019 |
| 4DNESL8HVDR5 | contact | list-combined | 5.52E+07 | 2.88E+07 | 1.16E+07 | 1.48E+07 | 11/4/2019 |
| 4DNES42T87S8 | contact | list-combined | 6.25E+07 | 3.19E+07 | 1.31E+07 | 1.75E+07 | 11/4/2019 |
| 4DNESDRECPY8 | contact | list-combined | 5.89E+07 | 2.87E+07 | 1.23E+07 | 1.79E+07 | 11/4/2019 |
| 4DNESITQCT9N | contact | list-combined | 5.31E+07 | 2.52E+07 | 1.26E+07 | 1.54E+07 | 11/4/2019 |
| 4DNESLBDF71X | contact | list-combined | 6.39E+07 | 2.72E+07 | 1.47E+07 | 2.20E+07 | 11/4/2019 |
| 4DNESNG6M4GI | contact | list-combined | 6.19E+07 | 2.84E+07 | 1.49E+07 | 1.86E+07 | 11/4/2019 |
| 4DNES1UQQCQC | contact | list-combined | 6.00E+07 | 2.47E+07 | 1.36E+07 | 2.17E+07 | 11/4/2019 |
| 4DNESXQ13LRW | contact | list-combined | 7.23E+07 | 3.21E+07 | 1.57E+07 | 2.45E+07 | 11/4/2019 |
| 4DNESW54C11P | contact | list-combined | 7.25E+07 | 2.97E+07 | 1.59E+07 | 2.70E+07 | 11/4/2019 |
| 4DNESOV38MUI | contact | list-combined | 7.46E+07 | 3.37E+07 | 1.72E+07 | 2.37E+07 | 11/4/2019 |
| 4DNESOUACCOP | contact | list-combined | 5.67E+07 | 2.36E+07 | 1.35E+07 | 1.96E+07 | 11/4/2019 |
| 4DNESJN4ZXIC | contact | list-combined | 7.46E+07 | 3.24E+07 | 1.80E+07 | 2.42E+07 | 11/4/2019 |
| 4DNESES8DB3N | contact | list-combined | 6.26E+07 | 2.54E+07 | 1.53E+07 | 2.20E+07 | 11/4/2019 |

| | | | | | | | |
|---|---|---|---|---|---|---|---|
| 4DNESFRB6NSI | contact | list-combined | 6.01E+07 | 2.54E+07 | 1.53E+07 | 1.94E+07 | 11/4/2019 |
| 4DNESO57HS3X | contact | list-combined | 5.97E+07 | 2.40E+07 | 1.43E+07 | 2.14E+07 | 11/4/2019 |
| 4DNESFKQBEV1 | contact | list-combined | 4.62E+07 | 1.71E+07 | 1.04E+07 | 1.86E+07 | 11/4/2019 |
| 4DNESPFYORMI | contact | list-combined | 3.34E+07 | 1.71E+07 | 7.30E+06 | 8.93E+06 | 11/4/2019 |
| 4DNES2AHLK4O | contact | list-combined | 3.36E+07 | 1.57E+07 | 7.75E+06 | 1.01E+07 | 11/4/2019 |
| 4DNES2R93GH1 | contact | list-combined | 4.96E+07 | 2.37E+07 | 1.10E+07 | 1.48E+07 | 11/4/2019 |
| 4DNES6VPZLD6 | contact | list-combined | 4.79E+07 | 2.18E+07 | 1.13E+07 | 1.48E+07 | 11/4/2019 |
| 4DNES8BXQHNL | contact | list-combined | 6.86E+07 | 3.29E+07 | 1.43E+07 | 2.14E+07 | 11/4/2019 |
| 4DNES3J3DFUS | contact | list-combined | 4.04E+07 | 1.84E+07 | 8.86E+06 | 1.32E+07 | 11/4/2019 |
| 4DNESXZV4GUI | contact | list-combined | 4.37E+07 | 2.09E+07 | 9.61E+06 | 1.32E+07 | 11/4/2019 |
| 4DNESAF3KM2R | contact | list-combined | 3.99E+07 | 1.75E+07 | 9.09E+06 | 1.33E+07 | 11/4/2019 |
| 4DNESZNNP25B | contact | list-combined | 5.79E+07 | 2.61E+07 | 1.16E+07 | 2.03E+07 | 11/4/2019 |
| 4DNESEX4VDTH | contact | list-combined | 4.66E+07 | 2.19E+07 | 1.15E+07 | 1.32E+07 | 11/4/2019 |
| 4DNES98NQ39I | contact | list-combined | 6.83E+07 | 2.88E+07 | 1.55E+07 | 2.40E+07 | 11/4/2019 |
| 4DNESMYI53QK | contact | list-combined | 3.70E+07 | 1.44E+07 | 9.40E+06 | 1.32E+07 | 11/4/2019 |
| 4DNES86MYL3E | contact | list-combined | 4.47E+07 | 1.87E+07 | 1.06E+07 | 1.53E+07 | 11/4/2019 |
| 4DNES33L8EEV | contact | list-combined | 4.80E+07 | 1.77E+07 | 1.16E+07 | 1.87E+07 | 11/4/2019 |
| 4DNESPWGWJYA | contact | list-combined | 4.77E+07 | 1.63E+07 | 1.27E+07 | 1.87E+07 | 11/4/2019 |
| 4DNESJ55821X | contact | list-combined | 4.42E+07 | 1.69E+07 | 1.11E+07 | 1.62E+07 | 11/4/2019 |
| 4DNESF7LJ88J | contact | list-combined | 5.57E+07 | 2.04E+07 | 1.32E+07 | 2.22E+07 | 11/4/2019 |
| 4DNESM7PB81R | contact | list-combined | 5.36E+07 | 2.03E+07 | 1.22E+07 | 2.11E+07 | 11/4/2019 |
| 4DNESCMQ9JOF | contact | list-combined | 3.13E+07 | 1.11E+07 | 6.67E+06 | 1.36E+07 | 11/4/2019 |
| 4DNESM14SDMG | contact | list-combined | 4.54E+07 | 1.61E+07 | 1.15E+07 | 1.78E+07 | 11/4/2019 |
| 4DNES5GB1X5P | contact | list-combined | 3.23E+07 | 1.15E+07 | 8.18E+06 | 1.26E+07 | 11/4/2019 |
| 4DNES2RQ6BDT | contact | list-combined | 3.13E+07 | 1.04E+07 | 8.10E+06 | 1.28E+07 | 11/4/2019 |
| 4DNES2R6PUEK | contact | list-combined | 2.93E+09 | 1.35E+09 | 6.63E+08 | 9.17E+08 | 3/14/2020 |
| 4DNES18BMU79 | contact | list-combined | 5.33E+08 | 2.57E+08 | 8.22E+07 | 1.94E+08 | 2019-01 |
| 4DNESH4UTRNL | contact | list-combined | 2.02E+09 | 8.70E+08 | 3.71E+08 | 7.83E+08 | 2019-01 |
| 4DNESNYBDSLY | contact | list-combined | 1.20E+09 | 5.11E+08 | 2.55E+08 | 4.36E+08 | 2019-01 |
| 4DNES54YB6TQ | contact | list-combined | 1.59E+09 | 6.47E+08 | 3.56E+08 | 5.83E+08 | 2019-01 |
| 4DNESRE7AK5U | contact | list-combined | 2.96E+08 | 1.77E+08 | 6.14E+07 | 5.73E+07 | 2019-01 |
| 4DNES425UDGS | contact | list-combined | 6.19E+08 | 2.88E+08 | 1.21E+08 | 2.10E+08 | 2019-01 |
| 4DNESEPDL6KY | contact | list-combined | 5.58E+08 | 2.11E+08 | 1.43E+08 | 2.04E+08 | 2019-01 |
| 4DNESZW7OOTL | contact | list-combined | 1.88E+09 | 1.06E+09 | 3.21E+08 | 5.07E+08 | 2019-09 |
| 4DNESAL82BWY | contact | list-combined | 2.28E+09 | 1.29E+09 | 5.05E+08 | 4.83E+08 | 2019-09 |
| 4DNESMU2MA2G | contact | list-combined | 2.32E+09 | 1.28E+09 | 4.20E+08 | 6.17E+08 | 2019-09 |
| 4DNES1ONB8TD | contact | list-combined | 2.44E+09 | 1.18E+09 | 4.94E+08 | 7.64E+08 | 2019-09 |
| 4DNES8IIWFGK | contact | list-combined | 2.27E+09 | 1.23E+09 | 4.93E+08 | 5.40E+08 | 2019-09 |
| 4DNES1INHSG7 | contact | list-combined | 2.25E+09 | 1.30E+09 | 5.05E+08 | 4.43E+08 | 2019-09 |
| 4DNESWNF3Y23 | contact | list-combined | 4.77E+08 | 2.22E+08 | 1.73E+08 | 8.20E+07 | 2019-12 |
| 4DNESWLWNWV8 | contact | list-combined | 4.00E+08 | 2.31E+08 | 1.28E+08 | 4.20E+07 | 2019-12 |

| | | | | | | | |
|---|---|---|---|---|---|---|---|
| 4DNESL3AW546 | contact | list-combined | 4.49E+08 | 2.55E+08 | 1.54E+08 | 4.02E+07 | 2019-12 |
| 4DNESTI1YC1H | contact | list-combined | 4.39E+08 | 2.79E+08 | 1.19E+08 | 4.02E+07 | 2019-12 |
| 4DNESXX38FO6 | contact | list-combined | 4.14E+08 | 2.86E+08 | 1.07E+08 | 2.12E+07 | 2019-12 |

**Supplemental Table S2**. Sequencing depth of Hi-C maps in the 4DNucleome database used in our analysis.

Published Hi-C used as metric comparison

GM12878 (GSE63525)
HCT-116 (GSE104334)
K562 (4DNESU95RUNO)
HMEC (GSE63525)
HUVEC (GSE63525)
HELA (GSE63525)
IMR90 (GSE63525)
K562 (GSE63525)
KBM7 (GSE63525)
NHEK (GSE63525)
HAP1 (GSE95014)
H9 (GSE105028)

**Supplemental Table S3**. Hi-C datasets used for benchmarking the Ultra-Res dataset.
